# Deep-sea megabenthos communities of the Eurasian Central Arctic are influenced by ice-cover and sea-ice algal falls

**DOI:** 10.1101/515015

**Authors:** Rybakova Elena, Kremenetskaia Antonina, Vedenin Andrey, Boetius Antje, Gebruk Andrey

## Abstract

Quantitative camera surveys of benthic megafauna were carried out during the expedition ARK-XXVII/3 to the Eastern Central Arctic basins with the research icebreaker *Polarstern* in summer 2012 (2 August-29 September). Nine transects were performed for the first time in deep-sea areas previously fully covered by ice, four of them in the Nansen Basin (3571-4066m) and five in the Amundsen Basin (4041-4384m). At seven of these stations benthic Agassiz trawls were taken near the camera tracks for species identification. The observed Arctic deep-sea megafauna was largely endemic. Several taxa showed a substantially greater depth or geographical range than previously assumed. Variations in the composition and structure of megabenthic communities were analysed and linked to several environmental variables, including state of the sea ice and phytodetritus supply to the seafloor. Three different types of communities were identified based on species dominating the biomass. Among these species were the actiniarian *Bathyphellia margaritacea* and the holothurians *Elpidia heckeri* and *Kolga hyalina*. Variations in megafaunal abundance were first of all related to the proximity to the marginal ice zone. Stations located closer to this zone were characterized by relatively high densities and biomass of *B. margaritacea* (mean 0.2-1.7 ind m^-2^; 0.2-1.5 g ww.m^-2^). The food supply was higher at these stations, as suggested by enhanced concentrations of pigments, organic carbon, bacterial cell abundances and porewater nutrients in the sediments. The fully ice-covered stations closer to the North Pole and partially under multi-year ice were characterized by lower concentrations of the same biogeochemical indicators for food supply. These stations nevertheless hosted relatively high density and biomass of the holothurians *E. heckeri* (mean 0.9-1.5 ind m^-2^; 0.3-0.4 g ww.m^-2^) or *K. hyalina* (mean 0.004-1.7 ind m^-2^; 0.01-3.5 g ww.m^-2^), which were observed to feed on large food falls of the sea-ice colonial diatom *Melosira arctica*. The link between the community structure of megafauna and the extent and condition of the Central Arctic sea-ice cover suggests that future climate changes may substantially affect deep ocean biodiversity.

## Introduction

Benthic megafauna comprises marine animals exceeding 0.5-1 cm in size visible on seafloor images. They play an important role in benthic ecosystems through active recycling of sedimented organic matter, bioturbation and foodweb linkages. Megabenthos is a dynamic component of deep-sea ecosystems able to react rapidly to environmental changes [1]. Observations of benthic megafauna in the Central Arctic Basins are rare because of technical difficulties of sampling in the remote deep-sea region covered by permanent ice. Traditional sampling methods such as trawling are challenged by the ice-cover: vessels cannot keep steady speed and course in the ice. Hence, most previous studies of Arctic megafauna communities were confined to marginal seas such as the Barents [2, 3], Kara [4], Laptev [5, 6], Greenland [7, 8] and Chuchki seas [9, 10, 11, 12], the Beaufort shelf [13], the Fram Strait and areas around Svalbard [14, 15, 16, 17, 18, 19, 20]. Sampling of Central Arctic benthic megafauna from depths exceeding 2000 m began in the late 19^th^ century during expeditions of *Fram*, the Swedish Spitsbergen expedition *Vøringen*, the Danish *Ingolf* Expedition, *Thor* and *Ermak* [21, 22, 23, 24, 25, 26, 27, 28, 29]. Further contributions were made by Soviet expeditions on board *Sadko* and *Fedor Litke* [30, 31, 32] and by the drifting ice stations SP2, SP3, SP4 and SP5 [33, 34, 32].

The first extensive photographic observation of Arctic deep-sea megafauna was conducted in the second half of the 20^th^ century [35]. Quantitative studies of the Central Arctic megafauna are few and they are focused on the Canada Basin [36, 37, 38]. More recent photographic and video surveys of Arctic deep-sea megafauna were conducted in the Canada Basin during the expeditions 2002–23 of the Canadian Coastguard icebreaker *Louis S. St. Laurent* in 2002 [39] and HLY05-02 of the US Coastguard icebreaker *Healy* in 2005 [40]. These investigations were based on a small number of ROV dives and video transects; the most commonly observed taxa were identified only to major taxonomic groups, and authors considered most data as qualitative. So far the only knowledge of temporal and spatial variation of Arctic deep-sea megafauna based on photo transects is confined to the HAUSGARTEN observatory in the Fram Strait, between Spitzbergen and Greenland [14].

A remarkable characteristic of the Central Arctic deep-sea megafauna north of 82°N is its very low density, as previously shown for the Canada Basin with 0.1–0.9 ind m^-2^ at depths 3816-3843 m [40]. In contrast, at the HAUSGARTEN observatory (78-80°N), the deep Atlantic gateway to the Central Arctic, densities of 30-40 ind m^-2^ were found at 1650 m, and lower values of approximately 10 ind m^-2^ at 3000 m depth [14]. These values are two orders of magnitude higher than those shown for the Canada Basin. Recent compilations of data on Arctic benthos confirm a strong decline in biomass from the outer shelves to the inner basins of the Arctic Ocean [13, 41]. Similar patterns were described for the Arctic deep-sea macrofauna. Kröncke [42, 43] and Kröncke et al. [44] suggested that the ice-covered Arctic Eurasian Basin is one of the most oligotrophic regions of the world ocean. The authors reported a low abundance (from 0 to 850 ind m^-2^) and biomass (from 0 to 83 g m^-2^) of macrofauna based on quantitative box-core samples at water depths 1018-4478 m [42]. In a more recent investigation, Degen et al. [45] combined data from modern field studies with published and unpublished data from the past 20 years and confirmed that the abundance, biomass and production of benthic macrofauna were the lowest under the full, multiyear ice-cover, but increased close to the productive marginal ice zone. The recent synthesis of Vedenin et al. [46] for macrofauna also showed substantial relationships of quantitative characteristics with water depth and sea ice cover, both influencing food input to the deep-sea.

All of this suggests that the energy flux via sedimentation of primary produced organic matter is the key factor determining the abundance and biomass of benthic communities in the deep Arctic Ocean, which belongs to the most oligotrophic seas globally [47, 48, 49, 50, 51, 52, 53]. Additional factors such as availability of the hard substrate, currents, proximity to shelves and evolutionary history of basins appear to control diversity and community composition [49, 54]. Wlodarska-Kowalczuk et al. [55] suggested that latitudinal differences in diversity may not primarily result from differences in primary production and subsequent organic matter fluxes to the seafloor. According to these authors low species diversity in the northern North-Atlantic is determined by the limited pool of local species, resulting from geophysical properties and history of the region. Bluhm et al. [41] showed that communities of the deep basins and the outer shelf share more than half of the taxa. According to Mironov et al. [56] the biogeographical history of the Arctic Ocean is characterized by processes of fauna emergence and submergence.

In the present paper we focused on the question of spatial patterns in benthic megafauna as a consequence of spatial dynamics in sea ice cover in the Central Arctic basin. During the past four decades, significant reductions of the sea-ice cover and thickness and extension of the melting season were observed, and they are expected to continue as a result of global warming and Arctic amplification [57, 58, 59, 60, 61, 62, 63]. The sea-ice retreat leads to increasing light availability and this in turn may increase primary production [64, 65]. However, primary production in the Arctic is also severely nutrient-limited, and nutrient availability may further decrease owing to increase of ocean stratification as a result of warming and ice melt [66, 67, 68]. Average estimates of primary production in the ice-covered Central Arctic are low, with values of 1 to 25 g C m^-2^ yr^-1^ [69, 66]. The ice algae can be a key food source in Arctic marine food webs [70, 71]. Hence sea-ice loss could lead to decreased ice algal production and sedimentation affecting the quantity and quality of food available to benthic communities. The deposition to the seafloor of large aggregations of the sea-ice diatom *Melosira* is known for Arctic shelves [72, 73] and was observed to occur in the deep, ice-covered basins as a consequence of rapid sea-ice melt in 2012 [71]. However the overall contribution of the ice diatoms to the nutrition of benthos remains unknown. Limited data on impact of the ice algae production on benthic communities exists for shelf and slope communities of the Chukchi, Barents and Bering Seas [74, 72, 73].

This study was conducted in summer 2012, during the record minimum of the sea ice cover in the Arctic Ocean [71]. The FS *Polarstern* Cruise ARK-XXVII/3 to the ice-covered Eastern Central basins explored the effect of the ice cover, ice algae and phytoplankton production on benthic communities in the region of 82-89°N and 30-130°E. Using the towed video and photographic platform Ocean Floor Observation System (OFOS), massive accumulations of live and degraded ice diatom algae were observed for the first time in the abyssal of the Arctic Ocean at the depth of ~ 4000 m [71]. It was found that some benthic animals, foremost holothurians and ophiuroids, actively feed on algal patches.

In the present study we investigated the key factors structuring the distribution of abyssal megafauna in the Central Arctic at that time, including variations in sea-ice cover and biogeochemical variables indicating food supply by phytodetritus sedimentation. Our main aims included revealing 1) the structure of megafauna communities in the ice-covered basins and effect of the marginal ice zone and 2) the effect of algal food falls on megafauna. Also, we compared the Eurasian basin megafauna community composition and structure with that of adjacent regions, and identified several new depth and geographical records for a number of taxa. The obtained data provide a baseline for future studies in the changing deep-sea Arctic Ocean.

## 2. Material and methods

### 2.1 Study area, photographic survey and sampling

Photographic surveys were carried out during the expedition ARK-XXVII/3 in summer 2012 (2 August-29 September) in the Nansen and Amundsen basins (Fig 1). The seafloor was photographed using a towed Ocean Floor Observation System (OFOS) [75]. Nine stations (one transect per station) were performed: four in the Nansen Basin between 83-84°N and 18-110°E at depths 3571-4066 m, and five in the Amundsen Basin between 83-89°N and 56-131°E at depths 4041-4384 m (Fig 1). Stations 1–5 were situated closer to the marginal ice zone (MIZ) and were characterized by first-year sea ice (FYI), whereas stations 7–9 were situated at some distance from the ice edge and were characterized by some multiyear ice (MYI) [71]. Station 6 was somewhat closer to the ice edge in 2012 than in any previous year. The total analysed area of the seafloor comprised 16190 m^2^, ranging from 206 m^2^ to 3379 m^2^ per transect. The length of transects varied from 210 m to 5500 m (Table 1). The marginal ice zone was defined as the zone showing 30-50% of the ice cover.

**Fig 1.**
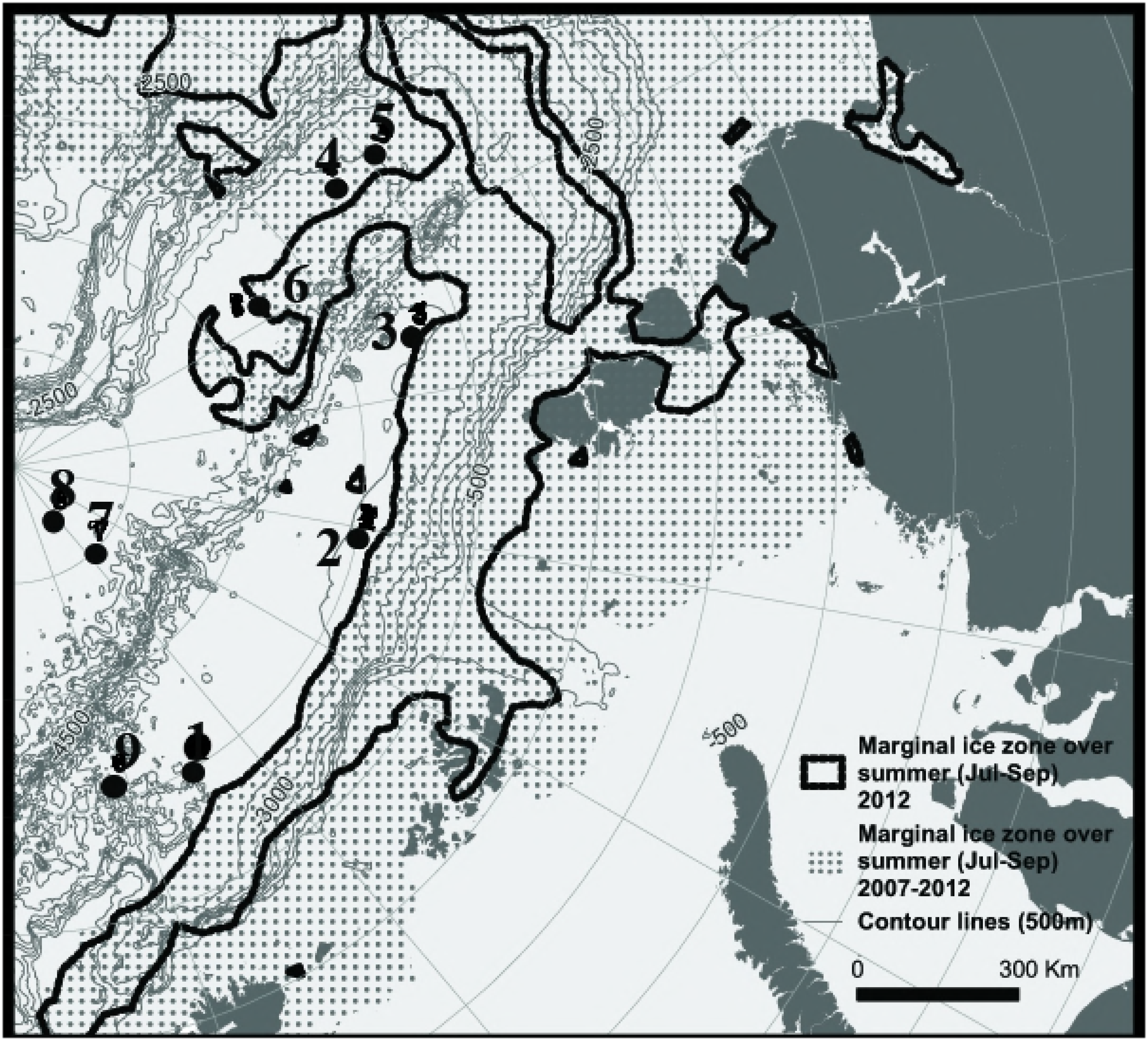
Location of stations performed during the expedition ARK-XXVII/3 in summer 2012 (August-September) to the Arctic Ocean. The ice margin in 2012 (black line) and the marginal ice zone integrated over previous 5 years (dotted area) where sea ice data were available are shown. 2012 represented a new sea-ice minimum except for an area around Gakkel Ridge (140-180°E) that showed a minimum already in 2007. Stations 1–5 were situated closer to the marginal ice zone and were characterized by the first-year sea ice. Stations 7–9 were situated at some distance from the ice edge and/or were characterized by multi-year ice (6).

**Table 1.**
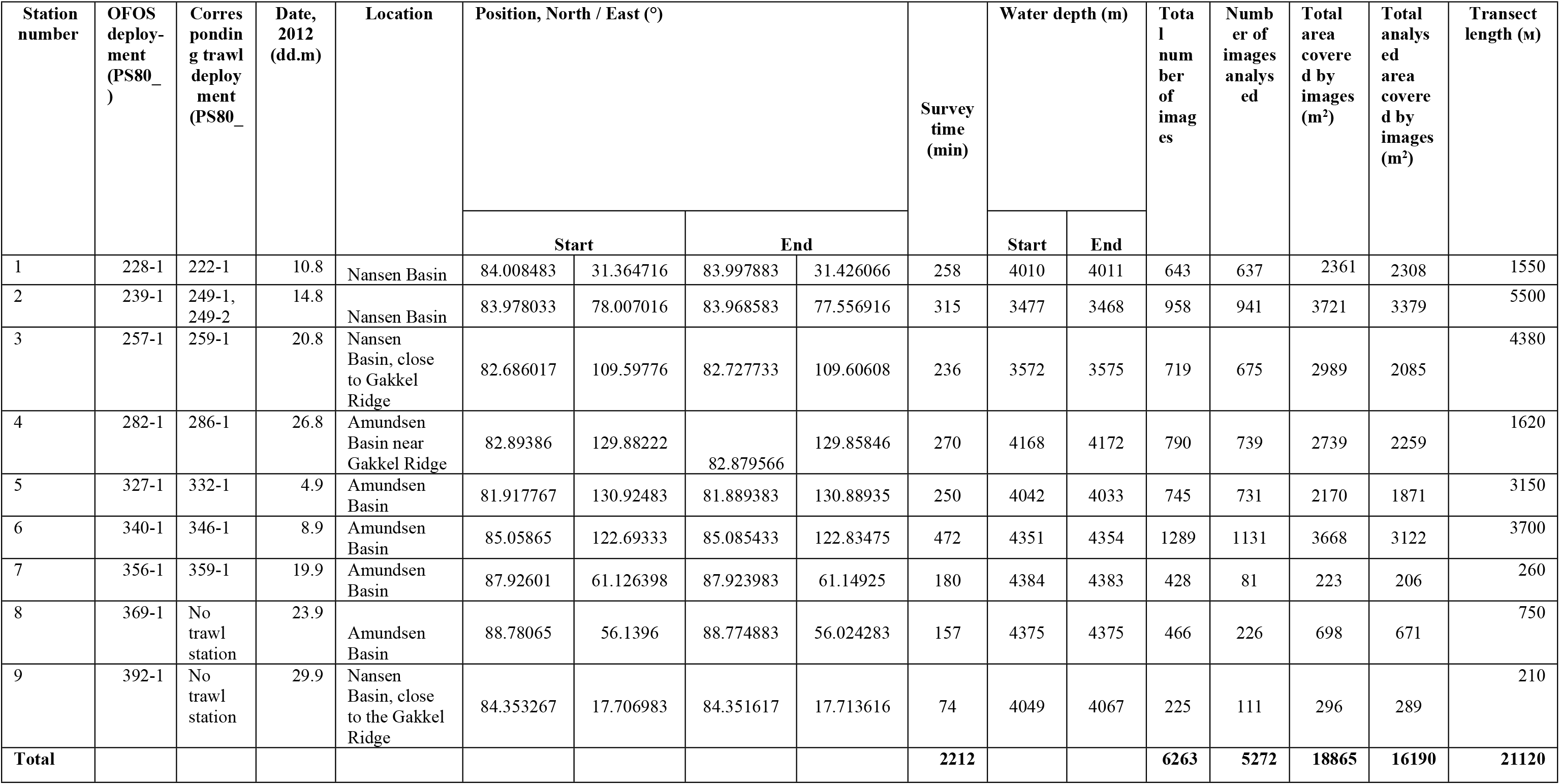
Details of OFOS transects.

The OFOS platform was equipped with the Canon camera EOS-1Ds Mark III (modified for underwater applications by iSiTEC GmbH, Germany), the strobe Kongsberg 0E11-242, four LED lights (LED Multi-Sealite, DeepSea Power and Light), telemetry (LRT-400 Fiber, iSiTEC) and three red laser points (OKTOPUS). A triangle laser scale with 50 cm between lasers was used to determine the camera’s footprint. The still camera was mounted to the frame in a vertical position to the seafloor. It was triggered automatically every 20 s, resulting in about 15-50 images per 100 m (for more details see Meyer et al. [75] and Soltwedel et al. [14]). Additional manually triggered images were taken when features of particular interest occurred in the field of view of the video camera, which was aligned with the still camera to help the winch operator adapting to bottom topography. These were however not used for quantitative statistical assessments. The OFOS was towed at approximately 0.1-1 knots in a ship drift for 1.5-8 hour of bottom time and at a targetaltitude of 1.3 m above the seafloor. This altitude has proven to be the optimal distance to the seafloor to achieve the best illumination and resolution of the images [14]. The area of seafloor on the images was in most cases 3–4 m^2^, of a total range of 1-9 m^2^ depending on target altitude. Start and end positions of transects (from GPS fixes) and water depths along transects (from echo soundings) were obtained from the ship’s data acquisition and management system DSHIP. Transect details, such as location, duration and the number of images are given in Table 1 and S1 Table.

Agassiz trawls were taken next to the photographic transects to obtain specimens for verification of taxonomic identifications based on images. In areas of stations 8 and 9 the thick ice prevented trawling operations. At stations 3, 4 and 7 the total trawled area (per station) was >1.5 times larger than the photographed area of the seafloor due to very low drift speed (<0.5 kn). At other stations the total trawled area was almost the same or two times smaller than the photographed area. Trawl samples were washed through a sieve with 1 mm mesh size, sorted and preserved in 4% buffered formaldehyde. Some specimens with calcareous skeleton were preserved in 80% ethanol for taxonomic identifications.

### 2.2 Image analysis, identification of megafauna and measurement of seafloor algal coverage

All images were analysed and stored using the image analysis program and database BIIGLE (Bio-Image Indexing, Graphic Labelling, and Exploration) web-2.0, which can be accessed by any standard web browser (www.BIIGLE.de) [15, 76, 77]. The laser points were used for calculation of the seafloor surface area on images. At transects 1-6 each image was treated as a separate sample, since there was no overlap between images. At transects 7-9 some overlap between images occurred owing to the low drift speed of around 0.1 kn, but overlapping images were excluded from the analysis. Images of unsatisfactory quality (with sediment clouds, too strong or low illumination, deviating distance from the bottom) were also excluded from the analysis. In total 6263 digital images were examined. Of these 5272 images were used for statistical analyses (Table 1).

All taxa and some habitat features were labelled in BIIGLE by species/feature name selected from a drop-down list [78]. Visible megafauna was counted and identified to the lowest possible taxonomic level. Taxonomic identifications were made with the assistance of zoological experts (see acknowledgements). The following taxa/organisms were excluded from statistical analyses: infauna represented only by surface traces, gelatinous zooplankton, small-size organisms (< 1 cm) and organisms that could not be identified at least to the phylum level.

The complete list of taxa identified on transects and in trawl samples is given in S2 Table.

The coverage of seafloor by algae aggregations and their remains was calculated based on sixty images at each transect using ImageJ software [79]. Images for this analysis were chosen with equal spatial intervals depending on the total number of images within a transect. The degree of freshness of algae in aggregations was evaluated visually using the two categories: greenish-brownish as freshly deposited, and whitish-yellowish as mostly degraded diatom falls.

### 2.3 Environmental parameters

Several environmental parameters were measured at the stations to assess possible effect on megabenthos communities. Characteristics of the sea ice included thickness, age and percentage of the ice cover. In the top 0-1 cm of sediment we analysed concentrations of chlorophyll a [Chl a] porosity (%), abundances of bacterial cells, total organic carbon (TOC), dissolved organic carbon (DOC), total dissolved nitrogen (TDN) and concentration of nutrients (PO_4_, Si, NO_2_, NO_3_). For more details of sediment characteristics measurements see Rossel et al. [53, 80]. All data from ARK-XXVII/3 were submitted to the Earth system database PANGAEA (see reference list for data collections [81, 82, 83, 84, 85, 86, 87].

### 2.4 Data analysis

The number of taxa was calculated at each station separately based on images and trawl samples. Based on images, asymptotic taxa richness values were estimated based on taxa accumulation curves using Chao 1 [88]. Chao 1 is influenced by the number of taxa that have only one or two individuals in the entire pool of samples. Permutated taxa-accumulation curves ware generated.

The quantitative analysis was performed only for data based on images; trawl data was used only for qualitative compositional studies. Trawl sampling is inefficient in catching some highly mobile epibenthic organisms, such as amphipods, isopods, shrimps and swimming polychaetes. Also usually there are difficulties with estimation of the seafloor area sampled by trawl. All taxa recognized on images were quantified for analyses with multivariate statistical methods and for estimations of biomass and density. The number of different taxa on each image was converted to individuals per m^2^ (density). Mean taxa densities (±standard deviation) and total megafauna density (±standard deviation) were calculated for each transect. The relative contributions (%) of the most abundant taxa to the total density were estimated.

The multivariate analysis was used to examine the structure of megafauna assemblages based on Bray-Curtis similarity coefficients and taxa density data. The Bray-Curtis similarity coefficient was calculated using square-root transformed data, owing to a large number of species represented by one individual and to reduce the dominance of the most abundant species. Non-metric multidimensional scaling plot (NMDS) was generated. Hierarchical cluster was overlain onto the NMDS plot. Analysis of similarity (ANOSIM) was used to assess the significance of differences among groups of stations revealed by NMDS. Contribution of taxa to similarity and dissimilarity of different groups of stations was calculated using the SIMPER subroutine of PRIMER v6 [88].

The following diversity indices were applied to describe megafauna assemblages: Pielou’s evenness (J’), Shannon–Wiener diversity (H’) (log_2_) and Simpson’s diversity (1-λ).

Spearman’s rank correlation was calculated between the environmental/sediment parameters and community characteristics or taxa densities. A canonical correspondence analysis (CCA) was performed at the station scale to estimate the contribution of environmental and sediment parameters to megafauna structure (DOC and TDN were not included because they were not measured at stations 1 and 5) [89]. Statistical analyses were performed in Primer V6 [90] and PAST V3 [91].

The mean taxa biomass per m^2^ and the total megafauna biomass per m^2^ were roughly estimated for each transect. The mean biomass (preserved wet weight) was calculated based on the wet weight of preserved individuals sampled by trawls. For taxa with insufficient trawl data the biomass was estimated using the biomass data of congeneric taxa or taxa with similar body shape; in some cases such taxa were excluded from analyses.

To examine the trophic community structure, nine most abundant taxa per transect were chosen and grouped according to their type of feeding. Feeding type was determined based on published data or original observations during the ARK-XXVII/3 cruise. Mean densities of organisms with different feeding types were calculated for all transects. The percentage of holothurians *Kolga hyalina* and *Elpidia heckeri* and the ophiuroid *Ophiostriatus striatus* associated with fresh and detrital algae aggregations was calculated for each transect. Biogeographic distribution patterns were evaluated for 43 species and 55 genera (all reliable identifications based on images and trawl samples). Seven generally accepted regions of the ocean were used as a reference in the biogeographical analysis: Arctic Ocean, North Atlantic, South Atlantic, Indian Ocean, North Pacific, South Pacific and Antarctic. Taxa present in all seven regions were classified as cosmopolitan. Endemic species are defined as those that were not reported outside the Arctic Ocean. The Greenland-Scotland Ridge separates the deep Arctic from Atlantic and shelves of the Chukchi and Bering Seas separate Arctic Basin from the Pacific [92].

## 3. Results

### 3.1 Taxonomic composition based on image data

In total 5272 images were analysed corresponding to the total area of 16190 m^2^ of the sea floor (Table 1). Overall 55000 specimens from 56 taxa were recognised, of which 52 taxa occurred in the Amundsen and 46 taxa in the Nansen Basin. Unique for either basin were 4 taxa in the Nansen and 14 in the Amundsen Basin. 30 taxa were identified to the species or genus level, 16 to the family; others were assigned to higher taxa. The number of taxa belonging to major taxonomical groups are presented in Table 2. The most species-rich group was the cnidarians with 16 taxa. Macellicephalinae accounted for the majority of polychaetes with at least eight taxa identified to the subfamily level. Lysianassoidea with five taxa (most of them at the family level) were the most diverse among amphipods. The number of taxa per transect varied from 19 (St. 7) to 43 (St. 6).

**Table 2.** Number of taxa in major taxonomic groups (image and trawl data).

| Taxon | Images | Trawl samples | Total |
| --- | --- | --- | --- |
| Porifera | 4 | 7 | 8 |
| Cnidaria | 16 | 0 | 3 |
| Ctenophora | 1 | 0 | 1 |
| Polychaeta | 11 | 9 | 17 |
| Echiura | 0 | 1 | 1 |
| Nemertea | 2 | 1 | 2 |
| Bivalvia | 1 | 4 | 4 |
| Gastropoda | 3 | 2 | 4 |
| Cephalopoda | 0 | 1 | 1 |
| Isopoda | 4 | 1 | 4 |
| Amphipoda | 11 | 7 | 15 |
| Mysida | 1 | 0 | 1 |
| Decapoda | 1 | 3 | 3 |
| Pycnogonida | 1 | 1 | 1 |
| Bryozoa | 1 | 2 | 2 |
| Ophiuroidea | 1 | 1 | 1 |
| Crinoidea | 1 | 1 | 1 |
| Holothuroidea | 2 | 2 | 2 |
| Asteroidea | 0 | 1 | 1 |
| Chaetognatha | 1 | 0 | 1 |
| Enteropneusta | 1 | 0 | 1 |
| **SUM** | **63** | **54** | **94** |

Many empty shells of shallow-water bivalves (*Astarte borealis, Mya truncata, Macoma* sp., *Cyrtodaria kurriana, Musculus* sp. and other unidentified) and gastropods (*Neptunea* sp. and other unidentified Neogastropoda) were noticed and marked on images. Their presence at abyssal depths remains unexplained, apparently they were transported by the transpolar drift of the sea-ice formed at sublittoral depths. Species represented only by empty shells were not included in the analyses.

Cumulative curves reached nearly asymptotic level at 2012 MIZ stations 2, 4 and 5 (Fig 2). Asymptotic taxa richness values (Chao 1) confirmed that a high fraction of predicted species richness was observed at these stations (Table 3). The value of Chao 1 was close to the observed number of taxa also at the multi-year ice stations 7 and 8, despite the relatively low number of analysed images (81 and 226 respectively). Thus we assume that the number of recorded taxa sufficiently documented real species composition at five stations, especially 2, 4 and 5. At stations 1, 3, 6 and 9 the species richness was undersampled as these stations comprised a high number of rare taxa (recorded only once or few times).

**Fig 2.**
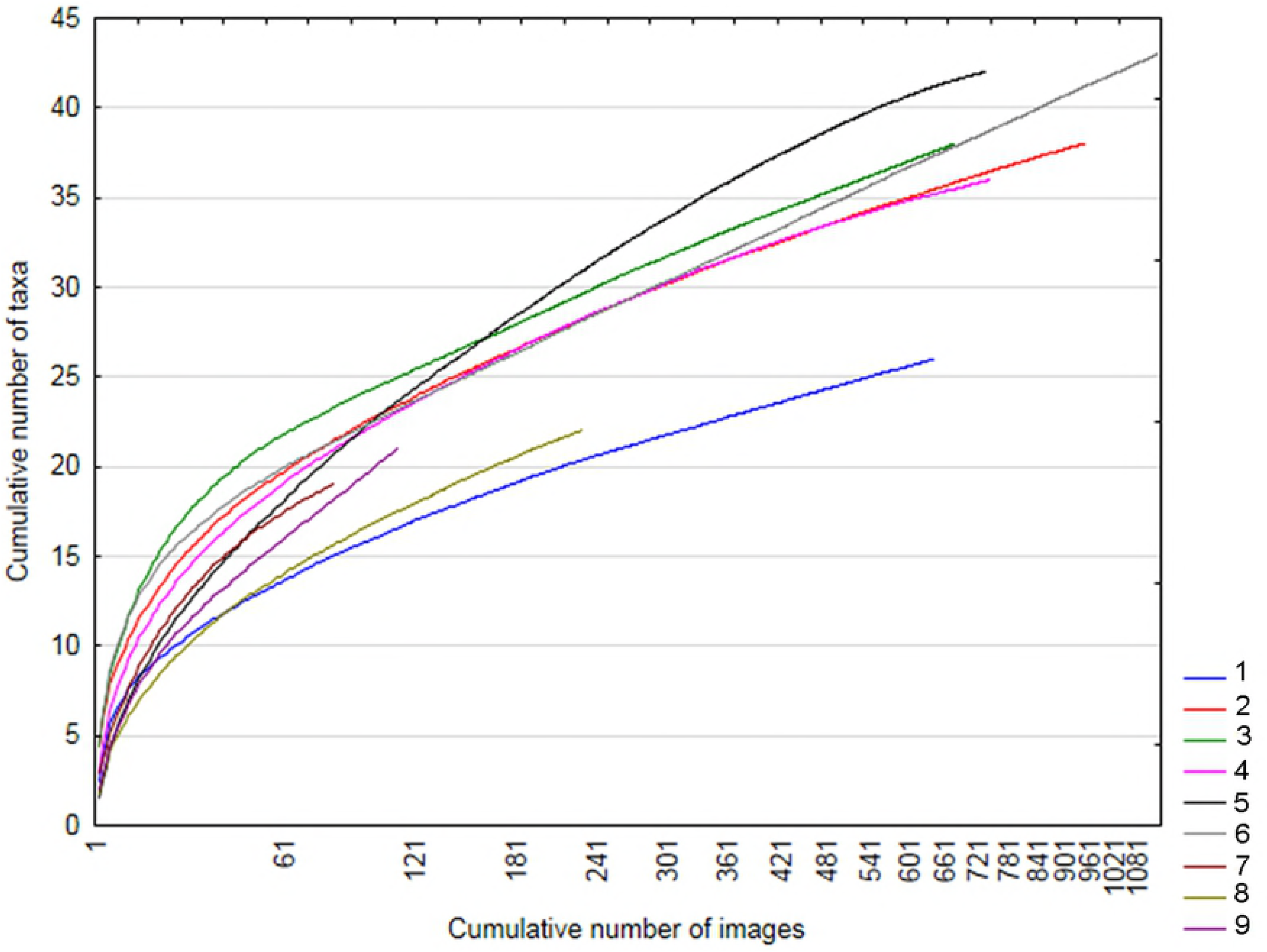
Image-based cumulative taxa-effort curves for nine stations. Effort is based on the cumulative number of images.

**Table 3.** Characteristics of benthic communities and algae aggregations on the seafloor. Total density, total biomass, observed (S) and estimated (Chao 1) taxa number based on images, species number in trawl samples, total number of taxa, Simpson diversity index(1-λ), Shannon diversity index (H’(loge)), Pielou’s evenness index (J’)) and seafloor algal coverage (%) and algal freshness are shown.

| Characteristic/Station | 1 | 2 | 3 | 4 | 5 | 6 | 7 | 8 | 9 |
| --- | --- | --- | --- | --- | --- | --- | --- | --- | --- |
| Total density (ind. m<sup>-2</sup>)<br>± SD | 1.1 ±0.6 | 2.93±1.2 | 3.3 ±1.3 | 7.8 ±5.3 | 3.7 ±1.9 | 4.2 ±1.5 | 3.7 ±1.2 | 0.9 ±0.6 | 1.5 ±0.7 |
| Total biomass (mg ww<br>m<sup>-2</sup>) | 240 | 1020 | 1640 | 770 | 1790 | 3940 | 600 | 210 | 350 |
| Observed taxa number<br>based on images (S) | 26 | 36 | 36 | 35 | 41 | 42 | 19 | 21 | 20 |
| Estimated taxa number<br>based on images<br>(Chao1) | 35 | 42 | 48 | 39 | 43 | 59 | 23 | 25 | 46 |
| Species number based<br>on trawl samples | 22 | 24 | 17 | 20 | 17 | 24 | 8 | - | - |
| Taxa number (Total) | 38 | 49 | 44 | 46 | 49 | 56 | 23 | 21 | 20 |
| Simpson diversity<br>index ($1-\lambda$) | 0.8 | 0.8 | 0.8 | 0.4 | 0.7 | 0.8 | 0.6 | 0.7 | 0.5 |
| Shannon diversity<br>index ($H'(\log e)$) | 1.8 | 1.9 | 2.0 | 1.0 | 1.8 | 1.8 | 1.2 | 1.7 | 1.2 |
| Pielou's evenness<br>index ($J'$) | 0.6 | 0.5 | 0.5 | 0.3 | 0.5 | 0.5 | 0.4 | 0.6 | 0.4 |
| Seafloor algal coverage<br>(%) ±SD | 0 | 0.03±0.04 | 1.6±0.4 | 0.3±0.2 | 0.5±0.2 | 0.4±0.6 | 2.4±0.7 | 10.3±0.8 | 0 |
| Algal freshness | absent | mostly fresh | mostly fresh, old<br>white degraded<br>patches present | mostly indistinct<br>old white<br>degraded patches | mostly not old<br>white degraded<br>patches | indistinct very<br>old big white<br>degraded<br>patches | mostly fresh, old<br>white degraded<br>patches present | a lot of small green<br>patches and big brown<br>patches, old white degraded<br>patches present | absent |

The six following taxa were abundant and common at all transects: the actiniarian *Bathyphellia margaritacea*, the serpulid polychaete *Hyalopomatus claparedii*, the polychaete Macellicephalin gen. sp.5, the isopod *Eurycope inermis* and the holothurians *Elpidia heckeri* and *Kolga hyalina* (Fig 3, S2 Table, S1 Figure). Eleven taxa were also common but less abundant, i.e. present on ~25% of images: the sponge Porifera gen. sp., the actiniarian Actiniaria gen. sp.1, unidentified animals in holes, the ceriantharian *Cerianthus* sp., the polychaete Macellicephalin gen. sp.1, the gastropod *Mohnia* sp., the amphipod *Eurythenes gryllus*, the lysianassid amphipod *Onisimus leucopis*, the mysid Mysidae gen. sp., the pycnogonid *Ascorhynchus abyssi* and the chaetognath *Pseudosagitta maxima*. Twelve taxa were referred to “rare” as they were noted only at one of the transects on a few images (S2 Table).

**Fig 3.**
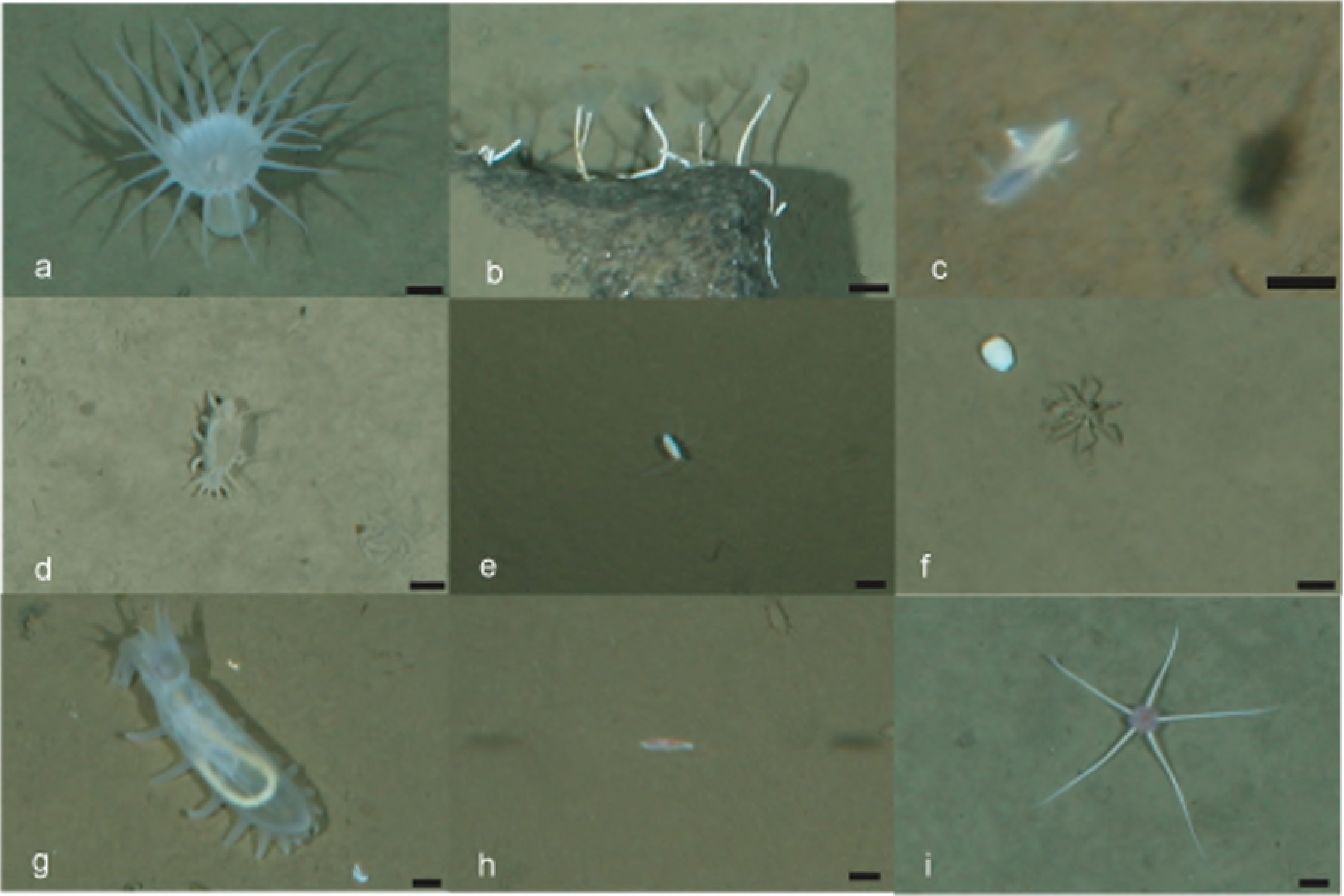
Images of the most abundant taxa. (a) *Bathyphellia margaritace* (Actiniaria), (b) *Hyalopomatus claparedii* (Polychaeta), (c) Macellicephalinae gen.sp.5 (Polychaeta), (d) *Elpidia heckeri* (Holothuroidea), (e) *Eurycope inermis* (Isopoda), (f) *Ascorhynchus abyssi* (Pycnogonida), (g) *Kolga hyalina* (Holothuroidea), (h) *Onisimus leucopis* (Amphipoda), (i) *Ophiostriatus striatus* (Ophiuroidea). Scale bar 1 cm.

### 3.2 Taxonomic composition based on trawl data

Trawl samples were taken at seven stations (MIZ stations 1-5, and fully ice-covered stations 6 and 7). In total 2131 specimens of benthic organisms were encountered in the trawl catches. The total megafauna density in trawl samples (0.1-0.6 ind m^-2^) was much lower when compared to values based on images (0.9-7.8 ind m^-2^). The most abundant epifaunal species recorded by camera (*Kolga hyalina, Elpidia heckeri, Bathyphellia margaritacea* and *Ophiostriatus striatus*) were quantitatively underrepresented in the trawl catches (first of all *B. margaritacea*). This was also the case with mobile organisms (such as amphipods, isopods and swimming polychaetes) caught in very small numbers, potentially due to their ability to escape trawls. At the same time the number of epifaunal hydrozoans, bryozoans and sponges, inhabitants of hard substrata such as stalks of the sponge *Caulophacus*, was underestimated based on images. Also the infaunal polychaetes caught in trawls were missing on images.

Altogether, the trawled specimens were assigned to 53 taxa (32 taxa in the Amundsen Basin and 40 taxa in the Nansen Basin). The overlap with taxa identified on images was 40%. Taxa numbers per trawl varied from 8 (St. 7) to 24 (St. 2). Most abundant in trawl samples were the actiniarian *Bathyphellia margaritacea*, the ophiuroid *Ophiostriatus striatus*, the holothurian *Kolga hyalina*, the polychaete *Anobothrus lauberi* and the bryozoan *Nolella* cf. *dilatata*. The list of identified taxa with abundance and biomass values at each station is given in S3 and S4 Tables correspondingly. Numerous fragments of dead sponges *Caulophacus arcticus* with epifaunal hydrozoans and bryozoans were registered at all stations. Most diverse taxa were polychaetes (9 families), amphipods (7 species, 5 of them of the superfamily Lysianassoidea) and sponges (6 families). The overall diversity of Porifera and Hydrozoa in trawl samples was higher in comparison with images. The diversity of isopods in trawl samples was lower (only 1 taxon) compared to images (4 taxa). Also Macellicephalin polychaetes were diverse and abundant on images, whereas in trawl samples only one species was found.

### 3.3 Combined trawl and image data, fauna characteristic

Altogether at least 89 taxa were recognized in the study area: 73 taxa in the Amundsen Basin and 72 in the Nansen Basin. 23 taxa were found both in trawl samples and on images (S2 Table). 30 species were registered only in trawls; 36 only on images. The total number of taxa per station based on combined image and trawl data varied from 20 (St. 9) to 56 (St. 6). The number of taxa belonging to major taxonomic groups registered on images and trawls is given in Table 2.

Species names were obtained for 42 species. The biogeographic analysis revealed that 23 (55%) among them are endemic to the Arctic Ocean (Table S5). Cosmopolitan deep-sea species make up 5%. At the genus level, 7% among 55 analysed genera are Arctic endemics (Table S6). About half of the species belong to the widespread genera, 48% among the latter are cosmopolitan and 14% occur in five regions outside the Arctic.

Our findings expanded the bathymetrical or geographical range of 9 taxa: the actiniarian *Oceanactis bursifera*, the polychaetes *Tubularia regalis, Bouillonia* sp. *Hyalopomatus claparedi* and Echiuridae fam indet., the decapod *Bythocaris curvirostris*, the bryozoan *Eucratea loricate*, the asteroid *Tylaster wyllei* and one enteropneust (for details see S7 Table).

### 3.4 Community structure based on visible taxa abundance

The estimated taxa richness (Chao 1) was the highest at the ice-covered station 6 in the vicinity of Gakkel Ridge and the lowest at the multiyear-ice station 7 close to the North Pole. Station 6 also had the highest megafauna densities along with the MIZ station 4, the lowest densities prevailed at the MIZ station 1 and at the ice-covered stations 8 and 9 (Table 3, S1 Table). There was no statistically significant correlation between estimated taxa richness and megafauna abundance. Individual taxa density with standard deviation is given in S8 Table.

Images of the most abundant taxa are presented in Figure 3. Percentage of dominance of individual taxa differed between stations (Fig 4, Table 4). Stations 1 and 8 were dominated by the polychaete Macellicephalinae gen. sp. 5 and the actiniarian *B. margaritacea*. Also stations 2 and 3 were highly similar. These stations were located at shallower depths than other stations (3468-3575 m) and they were dominated by the actiniarian *Bathyphellia margaritacea* and the polychaete Macellicephalinae gen. sp. The ophiuroid *Ophiostriatus striatus* was found only at three stations and was notably abundant at stations 2 and 3. The dominance of *B. margaritacea*, Macellicephalinae gen. sp. 5 and the polychaete *Hyalopomatus claparedii* in different proportions was also observed at stations 4 and 5. At stations 7 and 9 (60% similarity) the dominants were the holothurian *Elpidia heckeri* and the polychaete Macellicephalinae gen. sp.5. Station 6 was dominated by the holothurians *Kolga hyalina* and *E. heckeri*. The highest dominance of a single taxon was at the MIZ station 4 (78%) and at the ice-covered station 9 (71%). At all other stations the value was <47%. Accordingly, evenness was the lowest at station 4 and the highest at the MIZ station 1 and ice-covered station 8. Diversity was the lowest at stations 4 and 9 and the highest at the MIZ station 3 (Table 3).

**Fig 4.**
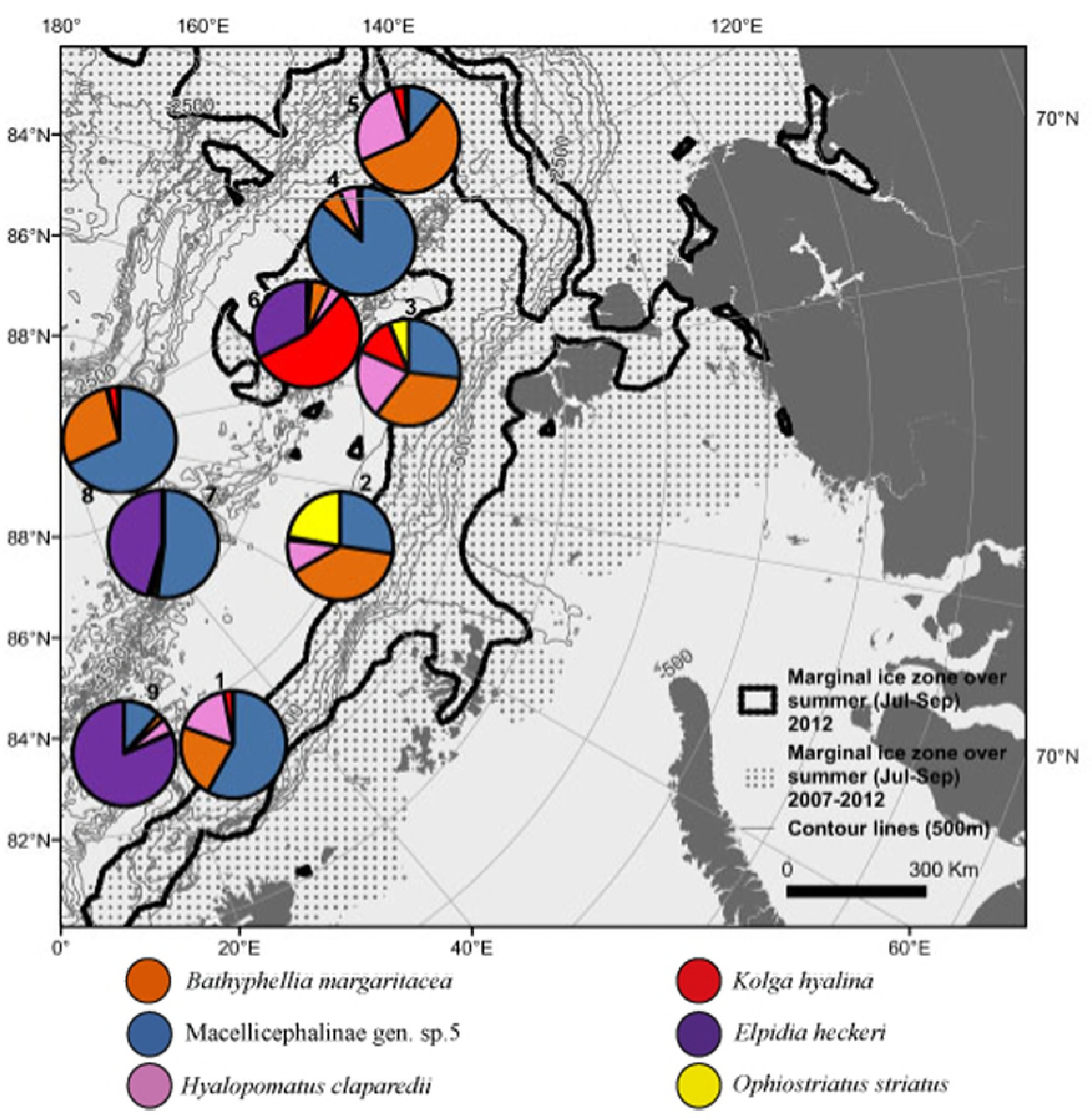
Contribution to abundance (in %) per station of six most abundant taxa.

**Table 4.**
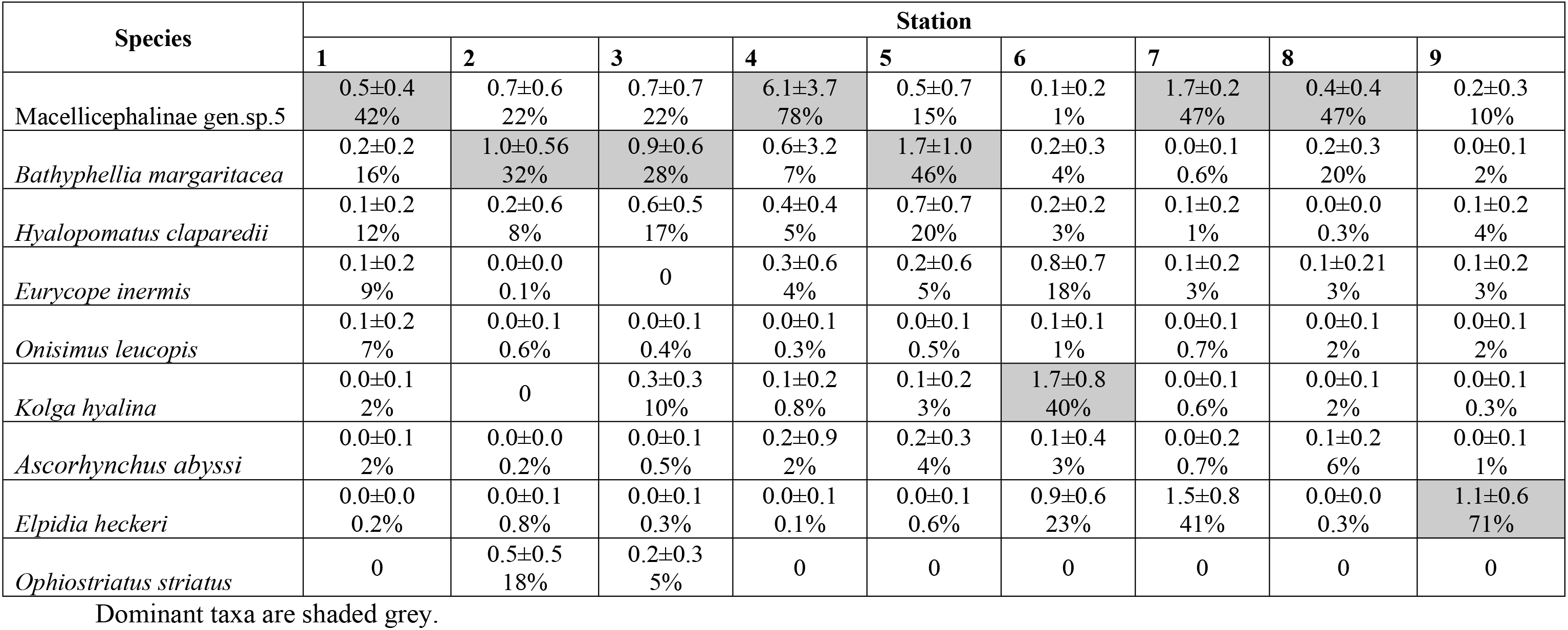
Mean density (ind. m^-2^) ± standard deviations and contribution to abundance (in %) of the most abundant taxa at nine stations.

| Species | Station |  |  |  |  |  |  |  |  |
| --- | --- | --- | --- | --- | --- | --- | --- | --- | --- |
|  | 1 | 2 | 3 | 4 | 5 | 6 | 7 | 8 | 9 |
| Macellicephalinae gen.sp.5 | 0.5±0.4<br>42% | 0.7±0.6<br>22% | 0.7±0.7<br>22% | 6.1±3.7<br>78% | 0.5±0.7<br>15% | 0.1±0.2<br>1% | 1.7±0.2<br>47% | 0.4±0.4<br>47% | 0.2±0.3<br>10% |
| *Bathypheilia margaritacea* | 0.2±0.2<br>16% | 1.0±0.56<br>32% | 0.9±0.6<br>28% | 0.6±3.2<br>7% | 1.7±1.0<br>46% | 0.2±0.3<br>4% | 0.0±0.1<br>0.6% | 0.2±0.3<br>20% | 0.0±0.1<br>2% |
| *Hyalopomatus claparedii* | 0.1±0.2<br>12% | 0.2±0.6<br>8% | 0.6±0.5<br>17% | 0.4±0.4<br>5% | 0.7±0.7<br>20% | 0.2±0.2<br>3% | 0.1±0.2<br>1% | 0.0±0.0<br>0.3% | 0.1±0.2<br>4% |
| *Eurycope inermis* | 0.1±0.2<br>9% | 0.0±0.0<br>0.1% | 0 | 0.3±0.6<br>4% | 0.2±0.6<br>5% | 0.8±0.7<br>18% | 0.1±0.2<br>3% | 0.1±0.21<br>3% | 0.1±0.2<br>3% |
| *Onisimus leucopis* | 0.1±0.2<br>7% | 0.0±0.1<br>0.6% | 0.0±0.1<br>0.4% | 0.0±0.1<br>0.3% | 0.0±0.1<br>0.5% | 0.1±0.1<br>1% | 0.0±0.1<br>0.7% | 0.0±0.1<br>2% | 0.0±0.1<br>2% |
| *Kolga hyalina* | 0.0±0.1<br>2% | 0 | 0.3±0.3<br>10% | 0.1±0.2<br>0.8% | 0.1±0.2<br>3% | 1.7±0.8<br>40% | 0.0±0.1<br>0.6% | 0.0±0.1<br>2% | 0.0±0.1<br>0.3% |
| *Ascorhynchus abyssi* | 0.0±0.1<br>2% | 0.0±0.0<br>0.2% | 0.0±0.1<br>0.5% | 0.2±0.9<br>2% | 0.2±0.3<br>4% | 0.1±0.4<br>3% | 0.0±0.2<br>0.7% | 0.1±0.2<br>6% | 0.0±0.1<br>1% |
| *Elpidia heckeri* | 0.0±0.0<br>0.2% | 0.0±0.1<br>0.8% | 0.0±0.1<br>0.3% | 0.0±0.1<br>0.1% | 0.0±0.1<br>0.6% | 0.9±0.6<br>23% | 1.5±0.8<br>41% | 0.0±0.0<br>0.3% | 1.1±0.6<br>71% |
| *Ophiostriatus striatus* | 0 | 0.5±0.5<br>18% | 0.2±0.3<br>5% | 0 | 0 | 0 | 0 | 0 | 0 |

Dominant taxa are shaded grey.

The multi-dimensional scaling of community structure based on taxa abundance data demonstrated considerable dissimilarity between all stations (Bray-Curtis similarity coefficient was around 40%) (Fig 5). Two groups of stations clustered together: three northernmost stations located under mixed first and multi-year ice (stations 6, 7 and 9; group A) and five stations of the marginal ice zone in first year ice (stations 1, 2, 3, 4 and 5; group B). The community structure of station 8 (the northern-most, under multi-year ice) resembled more the composition of MIZ stations. ANOSIM indicated significant differences between the groups A and B (R=0.955, p=0.001). Similarity percentages routine (SIMPER) revealed that the average similarity of the group A was based on the similar abundance of *Elpidia heckeri*, Macellicephalinae gen. sp. 5 and *Eurycope inermis*. Within group B abundance of three species contributed most to the similarity: *Bathyphellia margaritacea*, Macellicephalinae gen. sp. 5 and *Hyalopomatus claparedii*. Dissimilarities within both groups correlated with the depth of stations (3500 m depth range vs. >4000 m). Within the group A station 6 separated from stations 7 and 9. Within the group B there was also subdivision: stations 2 and 3 separated from stations 1, 4, 5 and 8. Dissimilarity between the two groups within the group A was driven mainly by *Kolga hyalina, E. inermis* and Macellicephalinae gen. sp. 5. Dissimilarity between the two groups within the group B was caused mostly by differences in the abundance of *Ophiostriatus striatus* and Macellicephalinae gen. sp. 5.

**Fig 5.**
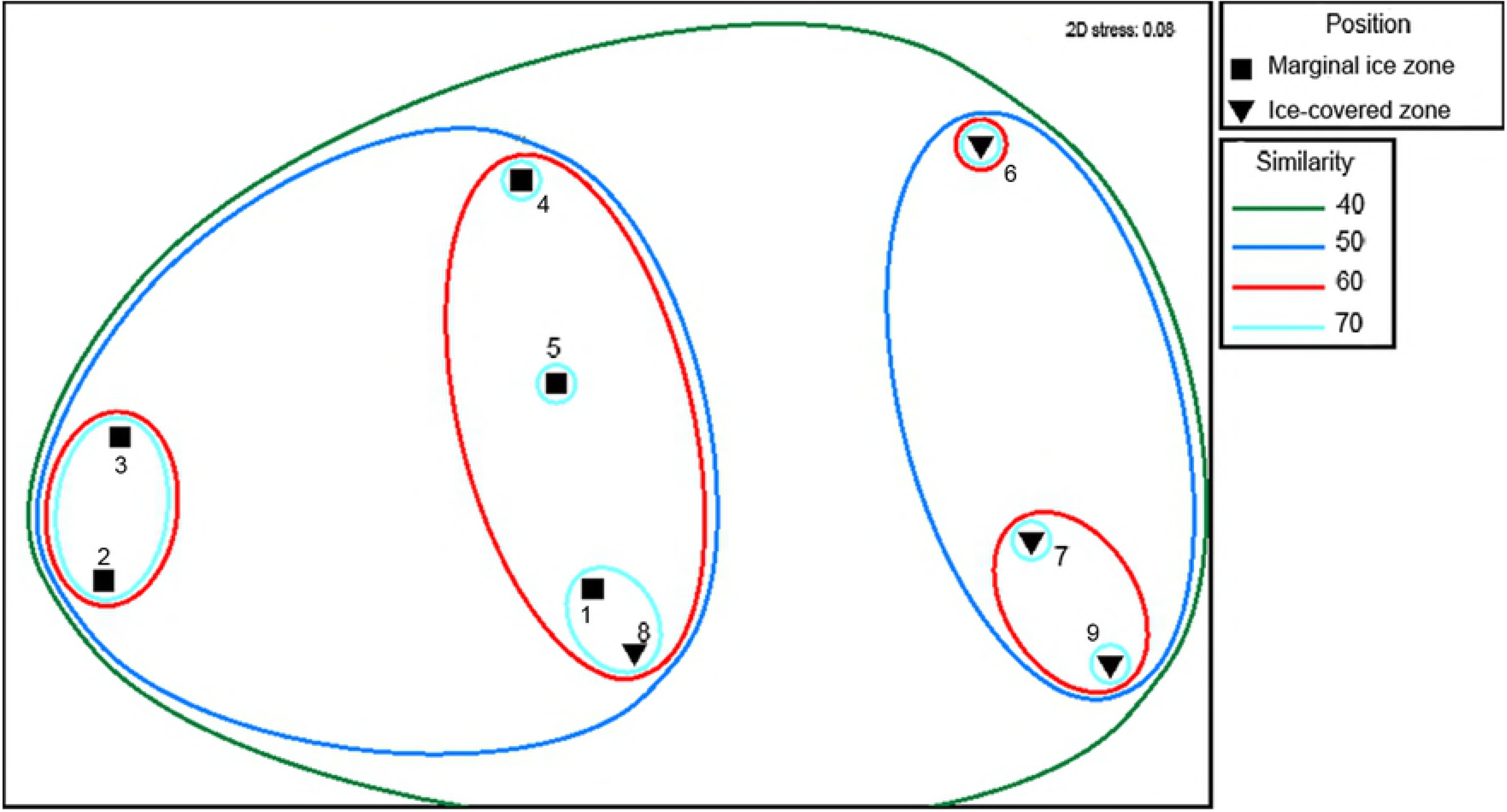
Multidimensional scaling plot for stations based on multivariate Bray-Curtis similarity coefficients for taxa abundance data. Abundance data were standardized to densities and square-root transformed. Similarity (%) is indicated by the coloured lines.

### 3.5. Biomass distribution

The highest megafauna biomass (based on estimations for image data) was at the ice-covered station 6 near Gakkel Ridge. High values were also obtained at the MIZ stations 2, 3 and 5. The lowest biomass was found at the ice-covered station 8 (Table 3, S1 Table).

Taxa dominating the biomass (nine in total) are shown in Fig 6 and Table 5. The actiniarian *Bathyphellia margaritacea* dominated at the MIZ stations 1, 2, 3, 4, 5 and at the ice-covered station 8. The holothurian *Kolga hyalina* was dominant at the ice-covered station 6. The holothurian *Elpidia heckeri* dominated at the ice-covered stations 7 and 9.

**Fig 6.**
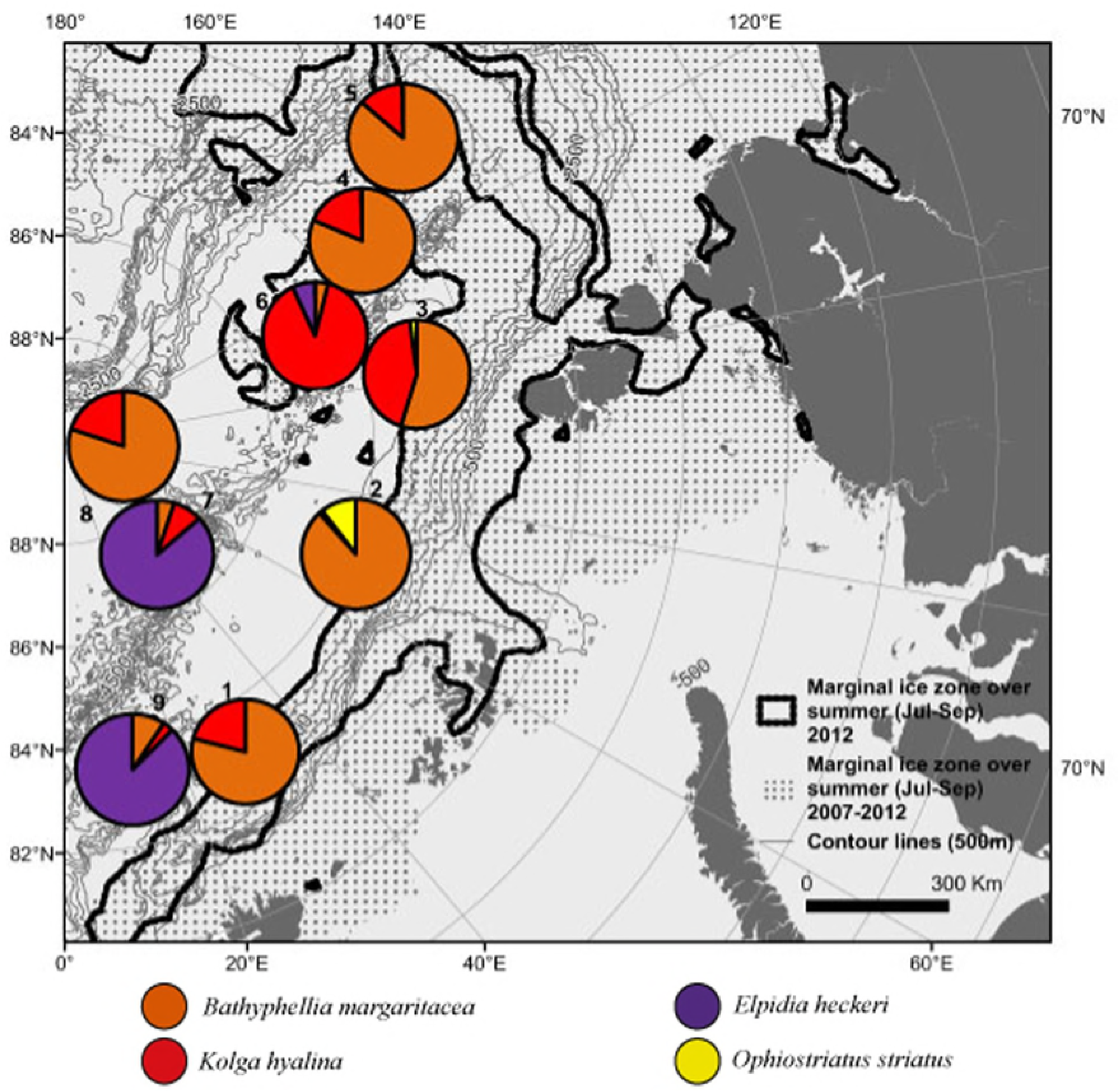
Contribution to biomass (in %) per station of four most abundant taxa.

**Table 5.**
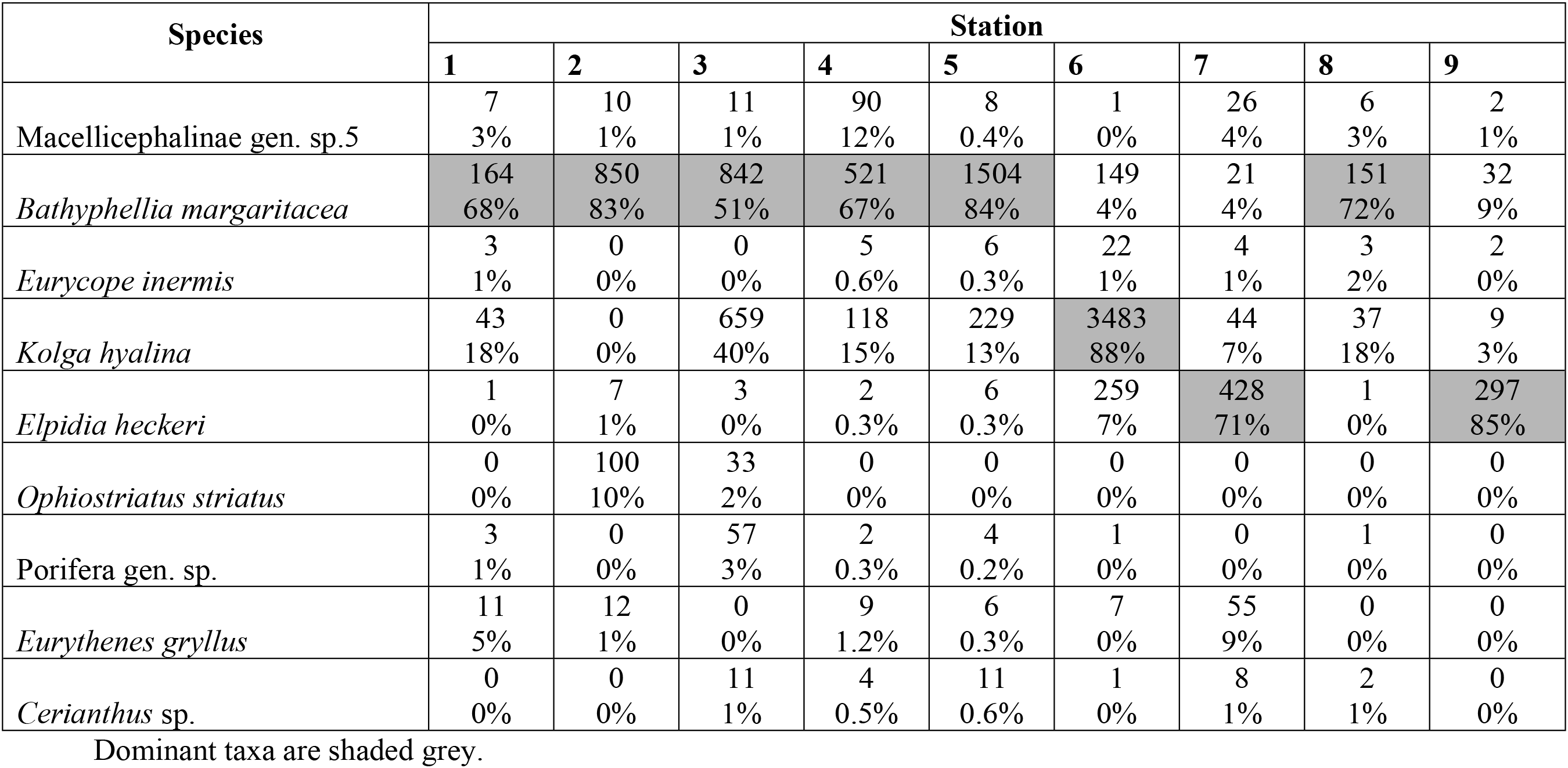
Mean biomass (mg ww m^-2^) and percent of total biomass of the most abundant taxa at nine stations.

| Species | Station |  |  |  |  |  |  |  |  |
| --- | --- | --- | --- | --- | --- | --- | --- | --- | --- |
|  | 1 | 2 | 3 | 4 | 5 | 6 | 7 | 8 | 9 |
| Macellicephalinae gen. sp.5 | 7<br>3% | 10<br>1% | 11<br>1% | 90<br>12% | 8<br>0.4% | 1<br>0% | 26<br>4% | 6<br>3% | 2<br>1% |
| *Bathypheilia margaritacea* | 164<br>68% | 850<br>83% | 842<br>51% | 521<br>67% | 1504<br>84% | 149<br>4% | 21<br>4% | 151<br>72% | 32<br>9% |
| *Eurycope inermis* | 3<br>1% | 0<br>0% | 0<br>0% | 5<br>0.6% | 6<br>0.3% | 22<br>1% | 4<br>1% | 3<br>2% | 2<br>0% |
| *Kolga hyalina* | 43<br>18% | 0<br>0% | 659<br>40% | 118<br>15% | 229<br>13% | 3483<br>88% | 44<br>7% | 37<br>18% | 9<br>3% |
| *Elpidia heckeri* | 1<br>0% | 7<br>1% | 3<br>0% | 2<br>0.3% | 6<br>0.3% | 259<br>7% | 428<br>71% | 1<br>0% | 297<br>85% |
| *Ophiostriatus striatus* | 0<br>0% | 100<br>10% | 33<br>2% | 0<br>0% | 0<br>0% | 0<br>0% | 0<br>0% | 0<br>0% | 0<br>0% |
| Porifera gen. sp. | 3<br>1% | 0<br>0% | 57<br>3% | 2<br>0.3% | 4<br>0.2% | 1<br>0% | 0<br>0% | 1<br>0% | 0<br>0% |
| *Eurythenes gryllus* | 11<br>5% | 12<br>1% | 0<br>0% | 9<br>1.2% | 6<br>0.3% | 7<br>0% | 55<br>9% | 0<br>0% | 0<br>0% |
| *Cerianthus* sp. | 0<br>0% | 0<br>0% | 11<br>1% | 4<br>0.5% | 11<br>0.6% | 1<br>0% | 8<br>1% | 2<br>1% | 0<br>0% |

Dominant taxa are shaded grey.

### 3.6 Aggregations of stalks of *Caulophacus arcticus*

Numerous aggregations of stalks of dead hexactinellid sponges *Caulophacus arcticus* were observed at stations 4, 5 and 6 (Fig 7a). Stalks of dead sponges sometimes co-occurred with «dark patches» of unknown origin (Fig 7b). At stations 4, 5 and 6, such aggregations were observed at 20-25% of images. The area of sponge aggregations varied from 0.02 to 1.3 m^2^ with the maximum coverage at St. 5. Stalks hosted numerous sessile fauna including actiniarians, hydroids, bryozoans and serpulid polychaetes, and also attracted mobile macellicephalin polychaetes, pycnogonids and isopods. Densities of sessile organisms and attracted fauna were up to 15-32 times higher in the presence of sponge stalks. For example, the density of actiniarians associated with dead sponge stalks increased to 9.7 ind m^2^, pycnogonids to 10.7 ind. m^2^, macellicephalin polychaetes to 6.9 ind. m^2^, isopods to 5.9 ind. m^2^, serpulid polychaetes to 2.8 ind. m^2^. Overall sponge stalk aggregations increased the local fauna density by 6 times (up to 30 ind. m^2^). Among those stations with aggregations of sponge stalks, the maximum faunal density was recorded at St.4 and was dominated by macellicephalin polychaetes. At St. 5 sponge stalks were mostly associated with the actiniarian *B. margaritacea*, and at St. 6 the holothurian *K. hyalina*.

**Fig 7.**
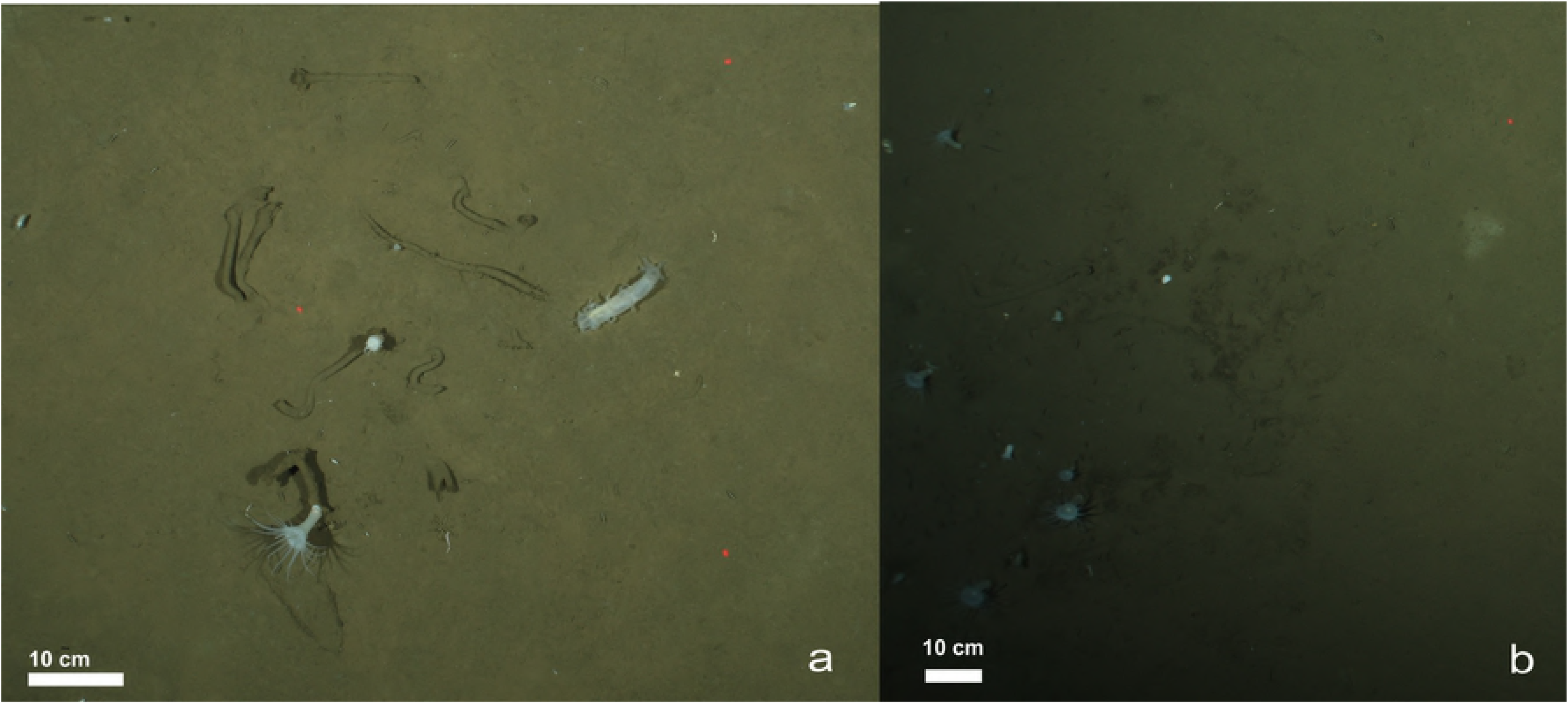
(a) Dead stalks of the hexactinellid sponge *Caulophacus arcticus* and associated fauna. (b) Accumulation of fauna at the sea floor near dead stalks of *C. arcticus* and undetermined «dark patches».

### 3.7 Algal coverage

The coverage of seafloor by algal patches varied from almost 0% (stations 1 and 9) to 10±1% (station 8) (Table 3, S1 Table). Patches varied from 5 to 12 cm^2^. At four stations (2,3,7 and 8) the algal falls were mostly fresh (based on visual parameters, Fig 8.). At other stations (4,5 and 6) mainly whitish remains of older algal aggregations were common. The oldest remains with no indications of fresh falls were observed at St. 6.

**Fig 8.**
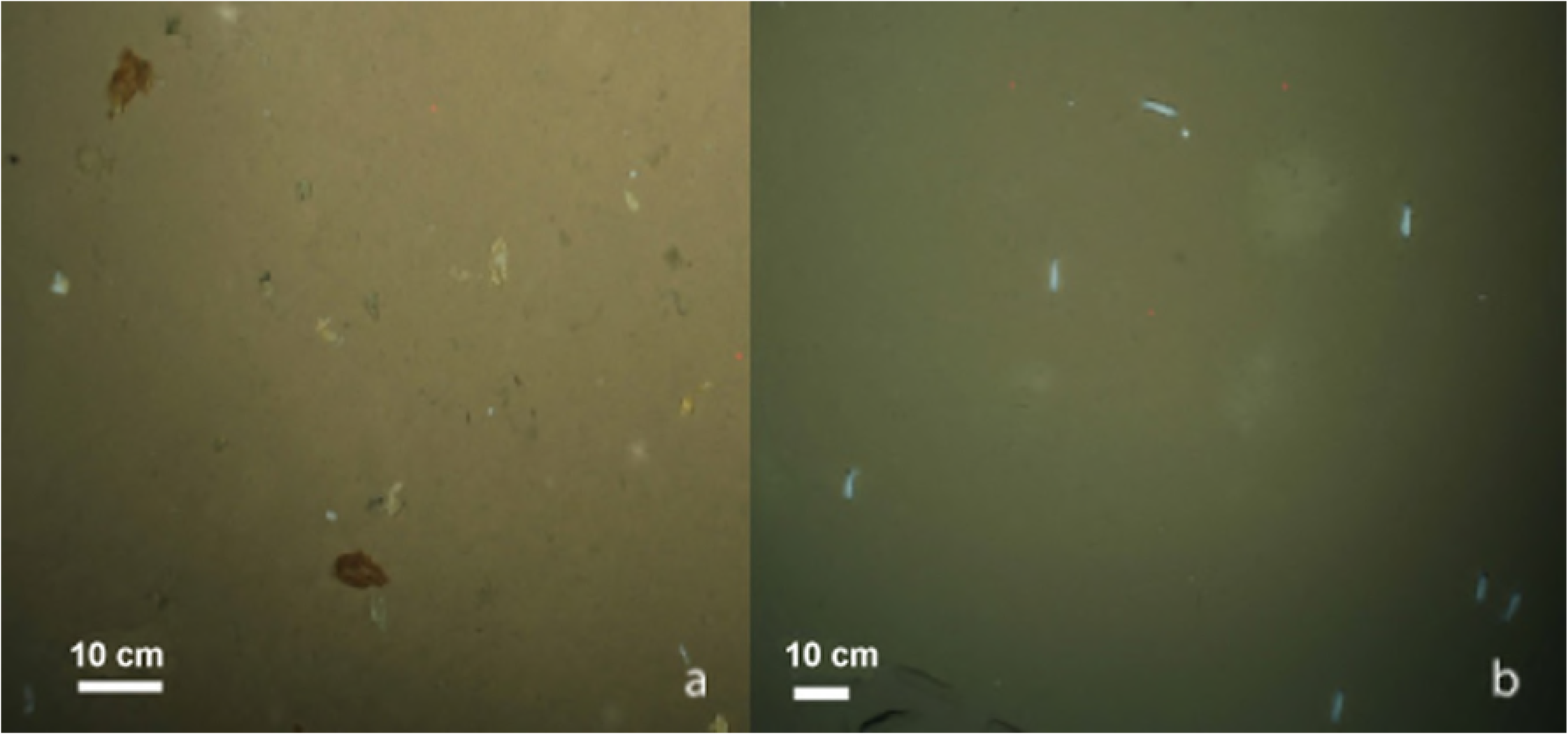
The degree of freshness of ice algae in aggregations at seafloor: (a) greenish-brownish, freshly deposited, (b) whitish-yellowish, mostly degraded diatom falls.

Three benthic species endemic to the Arctic were observed feeding on algae aggregations: the ophiuroid *Ophiostriatus striatus* and the holothurians *Kolga hyalina* and *Elpidia heckeri*. At stations with abundant *K. hyalina* (3,4,5 and 6), 10-40 % of specimens of this holothurian were associated with algal falls. Maximum values were recorded at stations with mostly fresh (St. 3) or moderately degraded (St. 5) algae. The percent of specimens of another holothurian, *E. heckeri*, associated with algae falls was similar: 10-44%, they dominated at St.7 with mostly fresh algae. The percentage of individuals of the ophiuroid *O. striatus* associated with algae falls was lower: about 5% at stations 2 and 3.

### 3.8 Feeding type

Suspension-feeders dominated densities (based on images) at all stations except for 6 and 9. More than 50% of individuals were *Bathyphellia margaritacea*, Macellicephalinae gen. sp. 5 and *Hyalopomatus claparedii* (Fig 9). Stations 6 and 9 were dominated by deposit-feeders (>65%) owing to high densities of the holothurians *Kolga hyalina* and *Elpidia heckeri*. The density of deposit feeders (first of all *E. heckeri*) also was high at St. 7 (45%). Density of carnivores/predator/scavengers (mainly *Ascorhynchus abyssi, Eurycope inermis* and *Onisimus leucopis*) was <25% at all stations.

**Fig 9.**
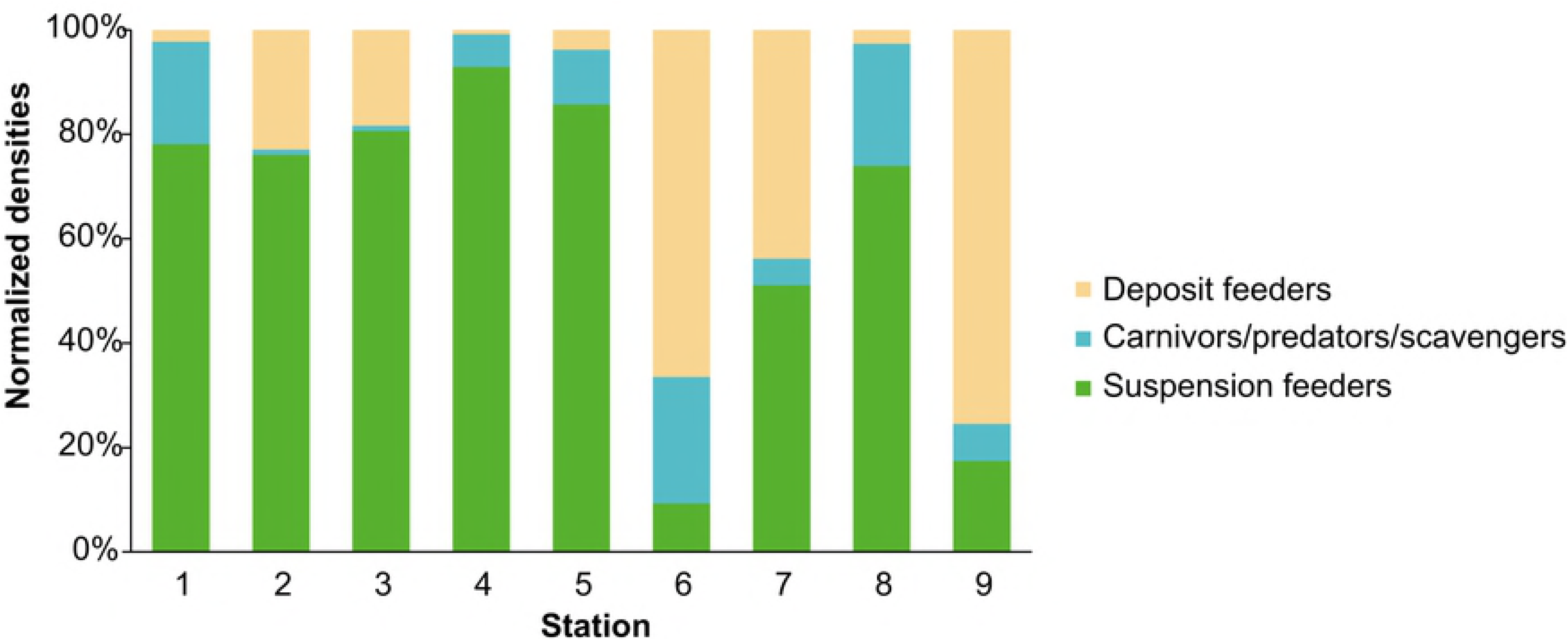
Mean densities of organisms with different feeding types (based on nine most abundant taxa at transects).

### 3.9 Relation between community structure and environmental parameters

Potential individual effects of single environmental factors on megafauna community composition and structure was evaluated using the observed variables presented in Table 6. Megafauna communities are known to depend on food supply, here indicated by concentrations of chlorophyll a extracted from sediments, as a proxy for the typical background phytodetritus sedimentation, and by the visually detected algal falls as a proxy for the specific 2012 melt-out of ice algae [71]. Additionally, bacterial cell numbers, dissolved organic carbon (DOC) and total organic carbon (TOC) concentrations were included in the analysis, indicative of longer term variations in food supply (multiyear to decadal time scale). Most of Spearman’s correlations between environmental parameters and integral community characteristics or individual species density/biomass were not statistically significant. Significant correlations are presented in Table 6. At stations with the ice coverage <50-80% (Stations 2, 3, 4, 5 and 6), the number of taxa was higher than at the northernmost stations (7, 8 and 9) with the ice coverage 100%, as demonstrated by cumulative curves (Fig 2). The estimated number of taxa (Chao 1) at northern stations 7 and 8 also was lower than at southern stations, and it was slightly different from the observed number of taxa at these stations. This suggests a negative correlation between richness of taxa and the sea ice coverage. The canonical correspondence analysis (Fig 10) revealed a cluster of stations with relatively high densities of the serpulid *Hyalopomatus claparedii* and the actiniarian *Bathyphellia margaritacea* (Stations 1,8,2,3 and 5). All of these stations except station 8 were associated with the ice margin and under the first-year ice. They were all characterized by relatively higher concentrations of chlorophyll a, TOC, abundance of bacterial cells, and a higher concentration of porewater nutrients. All of these variables showed lower concentrations at stations 6, 7 and 9 located at some distance from the ice edge, and at least partially under thicker multi-year ice and 100% ice cover. They showed a higher densities of the holothurians *Elpidia heckeri* (stations 9 and 7) and *Kolga hyalina* (station 6).

**Fig 10.**
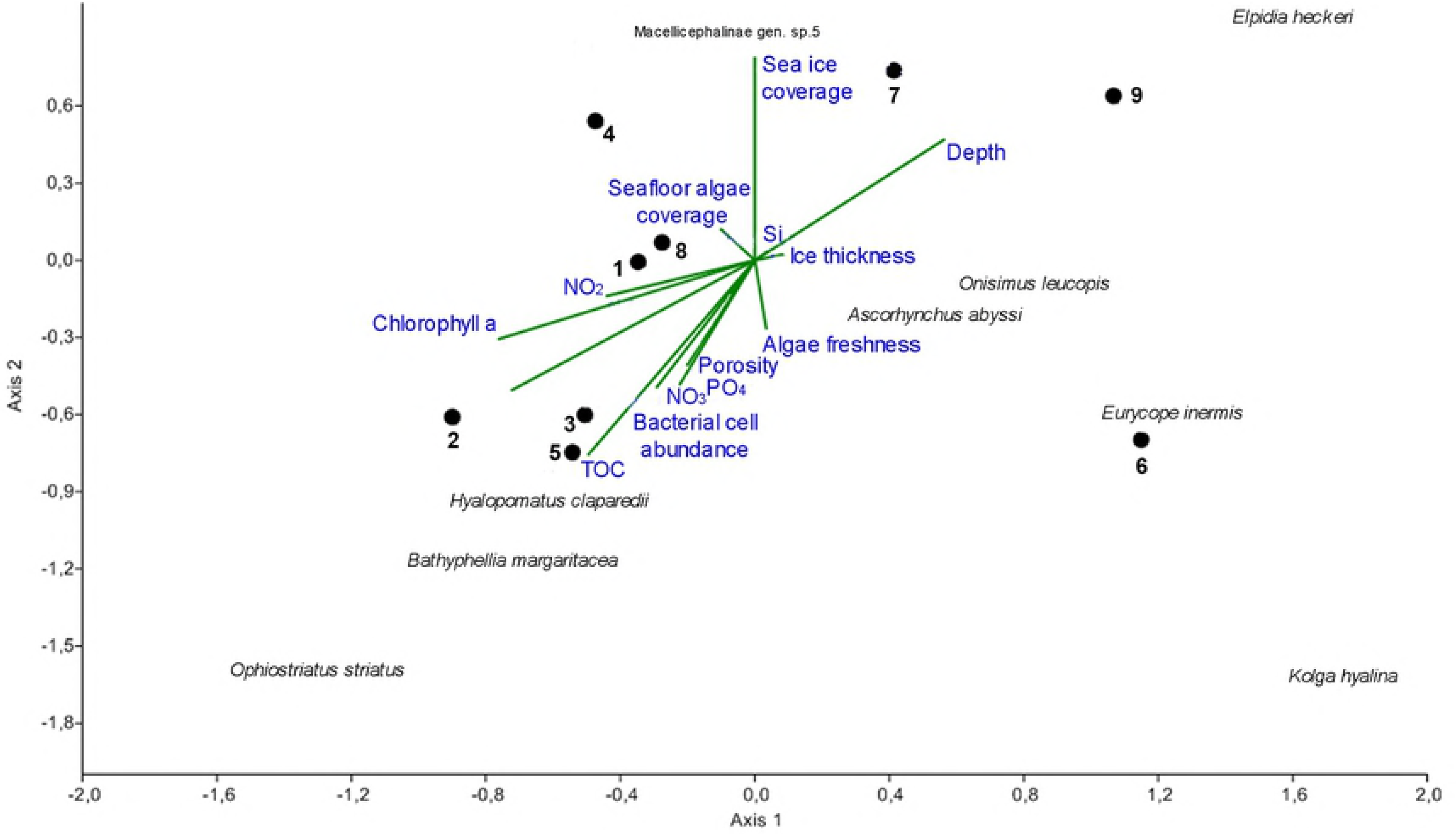
Canonical correspondence analysis (CCA). Explained variance on the first axis - 41% and second axis - 33%.

**Table 6.** Spearman’s rank correlation between the environmental/sediment parameters and community/species characteristics.

|  | **Number**<br>**of taxa** | *Bathypheilia*<br>*margaritacea*<br>(density) | *Hyalopomatus*<br>*claparedii*<br>(density) | *Eurycope*<br>*inermis*<br>(density) | *Onisimus*<br>*leucopis*<br>(density) | *Ascorhynchus*<br>*abyssi*<br>(density) | **Total**<br>**biomass** | *Bathypheilia*<br>*margaritacea*<br>(biomass) | *Cerianthus*<br>*sp.*<br>(biomass) |
| --- | --- | --- | --- | --- | --- | --- | --- | --- | --- |
| **Sea ice coverage** | -0,94<br>(0,0006) | p>0,05 | p>0,05 | p>0,05 | p>0,05 | p>0,05 | -0,73<br>(0,03) | p>0,05 | p>0,05 |
| **TOC** | 0,72<br>(0,03) | 0,9 (0,002) | 0,73 (0,03) | p>0,05 | p>0,05 | p>0,05 | p>0,05 | 0,9 (0,002) | p>0,05 |
| **Bacterial cell**<br>**abundance** | 0,74<br>(0,03) | 0,78 (0,01) | 0,77 (0,02) | p>0,05 | p>0,05 | p>0,05 | p>0,05 | 0,78 (0,01) | p>0,05 |
| **Chl a** | p>0,05 | 0,8 (0,01) | p>0,05 | p>0,05 | p>0,05 | p>0,05 | p>0,05 | 0,82 (0,01) | p>0,05 |
| **Depth** | p>0,05 | -0,82 (0,01) | p>0,05 | 0,78<br>(0,01) | p>0,05 | p>0,05 | p>0,05 | -0,82 (0,01) | p>0,05 |
| **First/multi year**<br>**ice** | p>0,05 | -0,87 (0,02) | -0,78 (0,03) | p>0,05 | p>0,05 | p>0,05 | p>0,05 | -0,87 (0,02) | p>0,05 |
| **DOC** | p>0,05 | p>0,05 | p>0,05 | -0,89<br>(0,01) | p>0,05 | -0,82 (0,02) | p>0,05 | p>0,05 | p>0,05 |
| **Algal freshness** | p>0,05 | p>0,05 | p>0,05 | p>0,05 | p>0,05 | 0,72 (0,04) | p>0,05 | p>0,05 | p>0,05 |
| **Seafloor algal**<br>**coverage** | p>0,05 | p>0,05 | p>0,05 | p>0,05 | p>0,05 | p>0,05 | p>0,05 | p>0,05 | 0,74 (0,03) |

Correlated pairs with $p < 0.05$ are shaded grey. Chl a – chlorophyll a, DOC - dissolved organic carbon, TOC – total organic carbon. The degree of algal freshness for correlation measurements was evaluated using categories: 0 – absent, 1 – mostly fresh, 2 – mostly old white degraded patches.

However, altogether, the effect of the algal coverage of the seafloor and freshness of algal aggregations was not statistically significantly related to total community structure and taxa abundance/biomass. None of the taxa (except for the ceriantharian *Cerianthus* sp.) showed a linear correlation with seafloor algal coverage. In fact, the lowest megafauna density (0.9 ± 0.6) was recorded at St. 8 with the maximum algal coverage (10%), despite the local aggregation of some *Elpidia heckeri* and *Kolga hyaline* feeding on the algal falls.

## 4. Discussion

This study aimed at providing quantitative data on the megafauna distribution and community structure in the ice-covered deep Nansen and Amundsen Basins of the Central Arctic. These basins show substantial change in sea ice cover due to warming, yet little is known how these changes are reflected in phytodetritus export and responses by the deep-sea benthos.

We were able to identify in this area 89 benthic taxa in total, based on images and trawl samples, and quantified their distribution to test the effect of ice cover and other spatial differences.

### 4.1 Taxa endemic to the Arctic Ocean

Endemicity at the species level was very high (55%) (<u>S5 Table</u>). Most of non-endemic species (24%) are shared with the deep North Atlantic. 19% of species are found both in the North Atlantic and North Pacific. At the genus level, the endemicity was low (7%) (S6 Table). Overall the examined fauna includes mainly the Arctic endemics or the Arctic-North Atlantic species belonging to genera widely distributed in the World Ocean. These results agree with trends shown by Mironov et al. (2013).

Among the Arctic endemics were all of the most dominant in our study species, such as *Bathyphellia margaritacea, Kolga hyalina, Elpidia heckeri* and *Onisimus leucopis*.

### 4.2. Sea ice and food supply structuring megafauna communities in the Eurasian basins

In earlier studies of pan-Arctic deep seas it was shown that the density and biomass of mega-, macro- and meiofauna increased in the marginal ice zone which is generally characterized by higher fluxes of phytodetritus compared to zones of permanent ice-cover [14, 15, 55, 93, 94]. The nine stations investigated in the present study have not been sampled before. They were separated by 52 – 689 nautical miles distance (Fig 1). The stations showed a substantial variation in the megafauna community structure, with a significant clustering of stations in relation to their proximity to the ice margin, which in 2012 (the year of study) had shifted northwards due to an unprecedented sea-ice minimum. The biomass at stations located closer to the ice margin (sts 1-5) was generally dominated by the actiniarian *Bathyphellia margaritacea*. In contrast, stations located further north in the fully ice-covered zone (sts 6, 7 and 9) were dominated by the holothurians *Elpidia heckeri* or *Kolga hyalina*. Though water depth in our study ranged only from 3400 m to 4400 m, we observed a biomass decrease with increasing depth fitting the globally predicted trend [95, 96]. However, most significantly, total biomass was negatively correlated with sea ice coverage and distance from the ice edge.

This substantial effect of ice-cover on community composition appears to be related mostly to variations in food supply. Several biogeochemical variables corresponding to the input of phytodetrital material on the seafloor demonstrate higher fluxes closer to the MIZ than at the northernmost stations. Because of the low bioturbation rate and relatively low degradation rate in polar waters, it is assumed that variables such as chlorophyll a pigments, total and dissolved organic carbon and nutrient concentrations reflect differences in food supply over the time scale of years to decades [47, 53]. The megabenthos biomass in the Nansen and Amundsen Basins in our study ranged from 0.21 to 3.94 g ww m^-2^, close to estimates of the macrobenthos biomass in the deep-sea Arctic [45, 46] and close to predicted calculated values for the Arctic fauna at depths 3500-4000 m [96].

Generally, both the biomass and density varied within an order of magnitude across stations, as did concentrations of biogeochemical indicators for food supply [71, 53]. However, the relationship between the composition and structure of megafauna communities and the observed biogeochemical indicators of food supply was not linear. A tendency of increasing abundance with increasing food supply was shown for the polychaete *Hyalopomatus claparedii* and the actiniarian *Bathyphellia margaritacea*. On the other hand, the abundance of holothurians *Elpidia heckeri* and *Kolga hyalina* declined with increasing food supply, both of which responded to the fresh food falls of sea ice algae. The dominant polychaete Macellicephalinae gen. sp. 5 showed no correlation with any parameters. The significance of correlations increased when the biomass values were used instead of abundance.

The present study also revealed a negative correlation between the number of taxa and the sea ice coverage for the Eurasian basins: the number of taxa was 17% lower at the northernmost stations under the multi-year ice compared to the MIZ stations under the first-year-ice. Earlier Wlodarska-Kowalczuk et al. [55] demonstrated that the species richness of the deep-sea Arctic fauna in general declines towards the North Pole. Authors concluded that the generally low species richness is preliminary determined by the limited pool of species available, resulting from the geophysical properties and history of the region. Our work indicates an important effect of ice-cover and food supply on diversity that needs further investigation, especially with regard to the effects of climate change in the Arctic.

### 4.3. Comparison with megafauna distribution in other Arctic regions

In the HAUSGARTEN area (the Fram Strait) unidentified anthozoans and the crinoid *Bathycrinus carpenteri* dominated the abundance at depths 2965-3043 m [14], i.e. slightly shallower than the depths dominated by the anthozoan *Bathyphellia margaritacea* in our study. Unidentified Actiniaria were among dominant taxa in the Canada Basin at depths 3843-3816 m [40]. The relatively high proportions of suspension feeding anthozoans at HAUSGARTEN, in Canada, Nansen and Amundsen Basins at depths 3000-4300 m is a characteristic pan-Arctic feature. Usually such suspension feeders are common at mid-slope depths [97].

In the present study unidentified swimming macellicephalin polychaetes were relatively abundant at depths 3570-4380 m. The dominance of unidentified swimming polychaetes was previously reported from the Canada Basin based on ROV observations (2800 m) [39]. This feature is unknown from Arctic shelves or deep-sea areas outside the Arctic Ocean.

Holothurians often dominate megafauna in the deep-sea [98], as also shown in our study. At the HAUSGARTEN observatory in the Fram Strait, *Elpidia heckeri* dominated the abundance at depths 5333-5404 m and *Kolga hyalina* at 2609-2629 m [14]. The first depth is somewhat deeper and the second shallower compared to our results. In the Porcupine Seabight in the North Atlantic the closely related to *K. hyalina* species *K. nana* was found in high densities (up to 50 ind m^-2^) between 2755 and 4080 m [99, 100].

Uneven distribution of holothurians at the abyssal seafloor, similar to our results, was demonstrated for the Canada Basin: a photographic survey revealed a high density of *Kolga hyalina* only at one of two stations, at 3816 m depth [40]. In another study in the Canada Basin by Bluhm et al. [39], during ROV observations only five holothurian specimens were recorded over 9.2 hours. The elpidiid holothurians are known to respond quickly to seasonal phytodetrital falls to the seafloor [101, 102, 103], which in part may explain patchiness of their distribution. The dominance of elpidiid holothurians in benthic communities is not limited to great depths. In earlier studies on the Celtic Margin (NE Atlantic) and in the north-western Canada Basin, high numbers of elpidiids were reported at much shallower depths: 1000–1500 m [104, 39].

The ophiuroid *Ophiostriatus striatus* was also numerous but not the dominant species at two shallower stations in the Nansen Basin. Ophiuroids often dominate the Arctic shelf epifauna reaching peak densities of several hundred ind. m^-2^ [72, 5, 8]. Bluhm et al. [39] reported high abundance of ophiuroids on the slope of the Northwind Abyssal Plain (1350 m). At HAUSGARTEN the ophiuroid *Ophiocten gracilis* occurred in high densities at the depth of approximately 1300 m [14].

### 4.3. Relationship between the ice algae aggregations on the seafloor and megafauna

Extensive aggregates of the sea-ice diatom *Melosira arctica* on the seafloor in the Arctic abyss were observed for the first time on the ARK-XXVII/3 expedition in 2012 [71]. This observation combined with the sediment trap data, benthic respiration rates and oxygen profiles in the sediment led to the hypothesis that extensive deposition of sea-ice algae to the deep seafloor was a consequence of rapid ice-melt in 2012, a new phenomenon for the deep-sea Arctic. Image analysis and investigation of the gut content of selected species showed that only some representatives of large mobile megafauna, such as the ophiuroid *Ophiostriatus striatus* and the holothurians *Kolga hyalina* and *Elpidia heckeri*, accumulated on ice-algae patches for feeding [71].

In the present study we quantified specific responses to and associations with these ice-algal falls. According to our image data, 40% of specimens of *Kolga hyalina* and 44% of *Elpidia heckeri* were associated with algal patches at the seafloor when the algal cover exceeded 0.5%. As these taxa dominated the megafauna in the same year when substantial algal falls occurred, we suggest that abundance of these species may increase significantly with reoccurring strong melt events, as observed in the current Arctic summers. However, we still found only weak evidence for a direct relationship between the megafauna abundance and biomass and the density of algal aggregations at the seafloor or the state of the algae (fresh vs. degraded). This supports the hypothesis that such algal food falls in the deep Arctic basins is a new phenomenon [71]. The biomass of only one species, *Cerianthus* sp., was positively correlated with the seafloor algae coverage. Also the density of only one species, the pycnogonid *Ascorhynchus abyssi*, was positively correlated with freshness of algae deposits.

Long-term studies in the North Atlantic and the North Pacific have previously demonstrated that abyssal megafauna is able to react rapidly to substantial changes in food supply. On the Porcupine Abyssal Plain in the North Atlantic two species of holothurians, *Amperima rosea* and *Ellipinion molle*, increased in abundance by more than two orders of magnitude suddenly in one year (1996) [101]. Following this event (known as the “Amperima event”), *A. rosea* was abundant over a wide area of the Porcupine Abyssal Plain indicating that this phenomenon was not a short-time fluctuation. Similar event occurred in the North Pacific in 2011 when densities of several elpidiid species sharply increased [103]. Also increases in holothurian abundance as a response to above-average localized food input were documented for *Elpidia glacialis* at Larsen A and B in Antarctica [105] and *Scotoplanes globosa*, two *Peniagone* sp. and an *Elpidia* sp. at Station M, 220 km west of the central Californian coast [106]. In our study the elpidiid *Kolga hyalina* occurred in the highest density at St. 6 with only old degraded algal deposits, indicating that the algal flux in that area took place before June [71]. Probably the high density of sea cucumbers at this station indicates that algal falls may have already occurred the year before and potentially provided energy for a higher population density.

In the Porcupine Seabight another species of this genus, *K. nana*, was reported to form very dense aggregations (up to 50 ind. m^-2^) in response to periodic fluxes of organic matter to the seafloor [99]. Based on our OFOS observations and observations in the Canada Basin [40] and the Porcupine Seabight [99], the species *K. hyalina* and *K. nana* are benthopelagic, able to swim in the near-bottom water layer [100]. This feature may help species of *Kolga* to respond rapidly to seasonal accumulations of organic matter at the seafloor. To fully understand the role of the ice-algae supply to the abyssal seafloor in the Arctic and the effect of this phenomenon on the abyssal benthic life, further studies are required.

### 4.7. Abyssal megafauna communities and climate change

Models predict continuing rapid warming of climate in the Northern Hemisphere causing the reduction of the sea-ice extent and thickness, including ice-free summers in the Arctic Ocean [59, 107]. Different scenarios of productivity changes are discussed, as not only the sunlight but also nutrient supply limits Arctic productivity and export flux [64, 65, 66, 67, 68]. Our study has demonstrated the relationship between the ice-cover, food supply and structure of deep-sea megafauna communities. Some species of deep-sea holothurians (for example *Amperima rosea*) react fast to algal deposits on the seafloor, and are able to selectively remove some biochemical compounds such as phytosterols from the freshly arrived phytodetritus in a short time of several months [108, 109]. This feature may have an impact on the food resource available to other deep-sea organisms.

Variations in the megabenthos community structure between stations in our study may reflect short and long term variations in phytodetritus flux to the seafloor. This flux is likely to depend on distance from the sea-ice margin and ice thickness. Hence, it can be expected that further warming and sea ice retreat will affect carbon flux to the abyssal and thereby the biodiversity and distribution of Arctic fauna. Unfortunately, no quantitative baseline data are available for the Central Arctic megabenthos before 2000. This challenges any assessment of temporal changes in the community structure. A decadal assessment of epibenthic megafauna of the HAUSGARTEN area in the Fram Strait over a period of 5 and 11 years showed significant changes in relative abundances of megafauna species that were related to variations in food supply with time, apparently linked to dynamics in sea ice cover and hydrography [15, 20].

The present study emphasizes the need for accumulating quantitative data on seasonal and annual variations of the Central Arctic megabenthos, to detect ecosystem changes in the deep-sea. The present work contributes to baseline data for the Nansen and Amundsen Basins, and to the first assessment of deep-sea benthos responses to sea-ice algae food falls.

## Conclusion

Our study on the composition and structure of megabenthos communities in different areas of the Eastern Central Arctic Basin combining quantitative photographic surveys with trawl sampling provides quantitative information of the dominant megafauna of the ice-covered basins and key factors structuring the distribution of abyssal megafauna in the Central Arctic. Three types of megafauna communities were distinguished: dominated by 1) the actiniarian *Bathyphellia margaritacea*, 2) the holothurian *Elpidia heckeri* and 3) the holothurian *Kolga hyalina*. Megafaunal abundance varied between stations, with some relation to the distance to the sea ice margin. Stations closer to the ice margin under first-year ice were characterized by relatively high densities and biomasses of *B. margaritacea* and relatively high food supply to the seafloor indicated by several biogeochemical variables. Stations located closer to the North Pole under the multi-year ice showed relatively low food supply, but relatively high densities and biomasses of holothurians *E. heckeri* and *K. hyalina* feeding on fresh algal falls of the colonial sea-ice diatom *Melosira arctica*. In case extensive algal food falls to the seafloor become regular due to increasingly frequent sea-ice melt events, the abundance of mobile deposit-feeding megafauna, such as elpidiid holothurians and ophiuroids, in the abyssal Central Arctic may rise significantly. Our data provide a baseline for future studies in changing deep-sea Arctic Ocean.

## Supporting information

**S1 Table. OFOS photographic survey transects characteristics, megafaunal communities characteristics, sea floor algae coverage and environmental conditions during POLARSTERN cruise PS80 (ARK-XXVII/3, IceArc).** (https://doi.pangaea.de/10.1594/PANGAEA.896626). (Pangaea, XLS)

**S2 Table. List of taxa in the OFOS photographic survey images and in Agassiz trawl samples during POLARSTERN cruise PS80 (ARK-XXVII/3, IceArc).** Frequency of occurrence of megafaunal taxa at OFOS transects is shown. (https://doi.pangaea.de/10.1594/PANGAEA.896618) (Pangaea, XLS)

**S3 Table. Abundance of megafauna collected by Agassiz trawl during POLARSTERN cruise PS80 (ARK-XXVII/3, IceArc).** (https://doi.pangaea.de/10.1594/PANGAEA.896627) (Pangaea, XLS)

**S4 Table. Abundance of megafauna collected by Agassiz trawl during POLARSTERN cruise PS80 (ARK-XXVII/3, IceArc).** (https://doi.pangaea.de/10.1594/PANGAEA.896629) (Pangaea, XLS)

**S5 Table. Characteristics of biogeographic distribution of species founded in the OFOS photographic survey and collected by Agassiz trawl during POLARSTERN cruise PS80 (ARK-XXVII/3, IceArc) to the Central Arctic Ocean in August and September 2012 (DOI)** (Supplementary, PDF).

**S6 Table. Characteristics of biogeographic distribution of genus founded in the OFOS photographic survey and collected by Agassiz trawl during POLARSTERN cruise PS80 (ARK-XXVII/3, IceArc) to the Central Arctic Ocean in August and September 2012 (DOI)** (Supplementary, PDF).

**S7 Table. New taxonomic findings and depth extension for megafauna founded in the OFOS photographic survey and collected by Agassiz trawl during POLARSTERN cruise PS80 (ARK-XXVII/3, IceArc) to the Central Arctic Ocean in August and September 2012.** Sample methods, location, depth and previously known distribution and depth range are shown (DOI) (Supplementary, PDF).

**S8 Table. Abundance of megafauna based on OFOS photographic survey during POLARSTERN cruise PS80 (ARK-XXVII/3, IceArc).**(https://doi.pangaea.de/10.1594/PANGAEA.896638) (Pangaea, XLS)

**S1 Figure. Catalogue of megafauna during POLARSTERN cruise ARK-XXVII/3** (https://www.pangaea.de/helpers/hs.php?s=Documentation&d=RybakovaE-etal_2018&t=Megafauna+during+POLARSTERN+cruise+ARK-XXVII/3&ID=896618) (Pangaea)

## Acknowledgements

We are grateful to the Captain, the officers and the crew of RV *Polarstern* for their support during the ARK XXVII/3 expedition. Our special thanks to experts for their help in taxonomic identifications: K. Tabachnik (Porifera), N. Sanamyan (Actiniaria), N. Budaeva (Polychaeta), A. Chernyshov (Nemertea), E. Krylova (Bivalvia), I. Nekhaev (Gastropoda), A. Raiskiy (Pycnogonida), G. Vinogradov (Amphipoda), M. Malyutina (Isopoda), A. Mironov (Echinoidea). We acknowledge Dr. Christina Bienhold for providing access to the environmental datasets. We also thank Dr. Vadim Mokievsky for informative discussions. Funding for this study was partly provided by the ERC advanced grant “Abyss” (Investigator grant no. 294757), and by the Alfred Wegener Institute (Program PACES II) to AB. The reported study was also partly funded by RFBR according to the research projects № 17-05-00787 to ER and № 18-05-60228 to AG and by the State assignment of IORAS (theme № 0149-2019-0009). This is publication no. awi-n#### of the Alfred Wegener Institute for Polar and Marine Research.

